# Pathological tau alters head direction signaling and induces spatial disorientation

**DOI:** 10.1101/2024.11.07.622548

**Authors:** Shan Jiang, Sara Hijazi, Barbara Sarkany, Verena G. Gautsch, Patrick A. LaChance, Michael E. Hasselmo, David Bannerman, Tim J. Viney

## Abstract

Spatial disorientation, an early symptom of dementia, is emerging as an early and reliable cognitive biomarker predicting future memory problems associated with Alzheimer’s disease, but the underlying neural mechanisms have yet to be fully defined. The anterodorsal thalamic nucleus (ADn) exhibits early and selective vulnerability to pathological misfolded forms of tau, a major hallmark of Alzheimer’s disease and ageing. The ADn contains a high density of head direction (HD) cells, which contribute to spatial navigation and orientation. Hence, their disruption may contribute to spatial disorientation. To test this, we virally expressed human mutant tau (htau) in the ADn of adult mice. HD-tau mice were defined by phosphorylated and oligomeric forms of htau in ADn somata and in axon terminals in postsynaptic target regions. Compared to controls, HD-tau mice exhibited increased looping behavior during spatial learning, and made a greater number of head turns during memory recall, indicative of spatial disorientation. Using *in vivo* extracellular recordings, we identified htau-expressing ADn cells and found a lower proportion of HD cells in the ADn from HD-tau mice, along with reduced directionality and altered burst firing. These findings provide evidence that expression of pathological human tau can alter HD signaling, leading to impairments in spatial orientation.

## Introduction

Spatial disorientation, which can be defined as a transient lack of awareness of one’s head direction with respect to spatial cues, is an early sign of dementia. Bouts of disorientation may occur prior to memory impairments (Liu et al., 1991; Pai and Jacobs, 2004; Serino and Riva, 2013; Costa et al., 2020; Colmant et al., 2022; da Costa et al., 2022). Therefore, disorientation has been suggested to be an early cognitive biomarker of dementia (Coughlan et al., 2018; Costa et al., 2020). However, the cause of these subtle behavioral changes remains unclear. The accumulation of abnormally hyperphosphorylated forms of tau (pathological forms of tau, ptau) has been shown to be associated with neurofibrillary degeneration and dementia. The Braak tau staging system has been widely adopted to classify the progression of ptau in Alzheimer’s disease (Braak et al., 2006), and the progression and extent of ptau in the cerebral cortex correlate with cognitive impairment including memory loss (Ossenkoppele et al., 2016; Bejanin et al., 2017; Harrison et al., 2019). We recently found that neurons in the human anterodorsal thalamic nucleus (ADn) accumulate ptau as early as Braak stage 0 (Sárkány et al., 2024), a stage when no cognitive impairments are expected. This nucleus was also found to exhibit extensive neurodegeneration in Alzheimer’s disease (Xuereb et al., 1991). In stark contrast, the adjacent anteroventral thalamic nucleus (AV) is unaffected at early Braak stages (Braak and Braak, 1991a; Sárkány et al., 2024).

The ADn contains a high density of head direction (HD) cells (Taube, 1995), which are thought to provide our sense of direction and play an important role in spatial navigation. HD cells are defined by an abrupt increase in firing when the head is oriented in a particular direction, i.e. their preferred firing direction (Taube et al., 1990). They are distributed across interconnected brain regions, including the lateral mammillary nucleus, anterior thalamus, dorsomedial striatum (DMS), retrosplenial cortex, postsubiculum (PoS), and entorhinal cortex (Taube et al., 1990; Wiener, 1993; Chen et al., 1994; Taube, 1995; Stackman and Taube, 1998; Mizumori et al., 2000; Sargolini et al., 2006; Taube, 2007; Clark et al., 2024). This HD network is essential for spatial orientation and navigation alongside grid cells, border cells, place cells and other kinds of spatially modulated cells forming a cognitive map (Gibson et al., 2013; Moser et al., 2017; Peyrache et al., 2017; Alexander et al., 2023). As a major node within this HD network, the ADn projects to other brain areas that are also enriched in HD cells, including the granular retrosplenial cortex (RSg) and PoS (Sripanidkulchai and Wyss, 1986; Van Groen and Wyss, 1990; Shibata, 1993b, a; Van Groen and Wyss, 1995; Cho and Sharp, 2001; Taube, 2007). The ADn is traditionally grouped with the AV and anteromedial thalamic nuclei forming the anterior thalamic nuclear group (ATN). Due to this grouping, lesion studies have largely overlooked the specific contributions of distinct nuclei. It is well established that ATN lesions in rodents cause spatial memory deficits (Aggleton et al., 1991; Aggleton et al., 1996; Warburton et al., 1997; Frost et al., 2021; Aggleton and O’Mara, 2022). Only recently, with the development of more specific targeting techniques, selective inhibition of the ADn alone has been shown to impair spatial working memory and alter contextual fear memory (Roy et al., 2021; Vetere et al., 2021; Roy et al., 2022), which is probably due to disruption of HD signaling. This raises the question as to whether selective accumulation of ptau reported in the human ADn at early stages could be detrimental to the functioning of HD cells (Shine et al., 2016).

To model the early accumulation of ptau within the HD network, we virally expressed a mutant form of human tau in the mouse ADn (HD-tau mice), leading to phosphorylated and oligomeric forms of human tau (htau) accumulating within cell bodies, dendrites and axons of ADn cells. We found that these HD-tau mice exhibited disorientation during spatial learning and recall, and HD cells had altered firing patterns, indicating that the accumulation of htau disrupts the HD network at early Braak stages, explaining the symptom of disorientation. Thus, HD-tau mice offer an opportunity to cross-validate an early cognitive biomarker for dementia, potentially leading to earlier detection and improved intervention strategies.

## Results

### Mapping the distribution of htau in the head direction network

To model early stages of ptau accumulation in the HD network, we bilaterally targeted the ADn of adult C57Bl6j mice with *AAV-CBh>EGFP:T2A:Tau(P301L):WPRE* or a control viral vector encoding EGFP only (Fig. 1A). Mutant (P301L) tau is known to misfold and aggregate under both viral and transgenic forms of expression (Lewis et al., 2000; Wegmann et al., 2019; Tetlow et al., 2023). After at least 8 weeks of expression, we mapped GFP fluorescence and htau immunoreactivity in the HD network (Fig. 1, S1; Table 1). HD-tau and HD-gfp (control) mice were defined by GFP+ somata predominantly localized to the ADn (Fig. 1A-D, S1A) and GFP+ axon terminals in postsynaptic target regions of the ADn, such as the thalamic reticular nucleus (TRN), DMS, RSg, and PoS (Fig. 1E, S1C; Table 1) (Shibata, 1993a, b).

**Figure 1.**
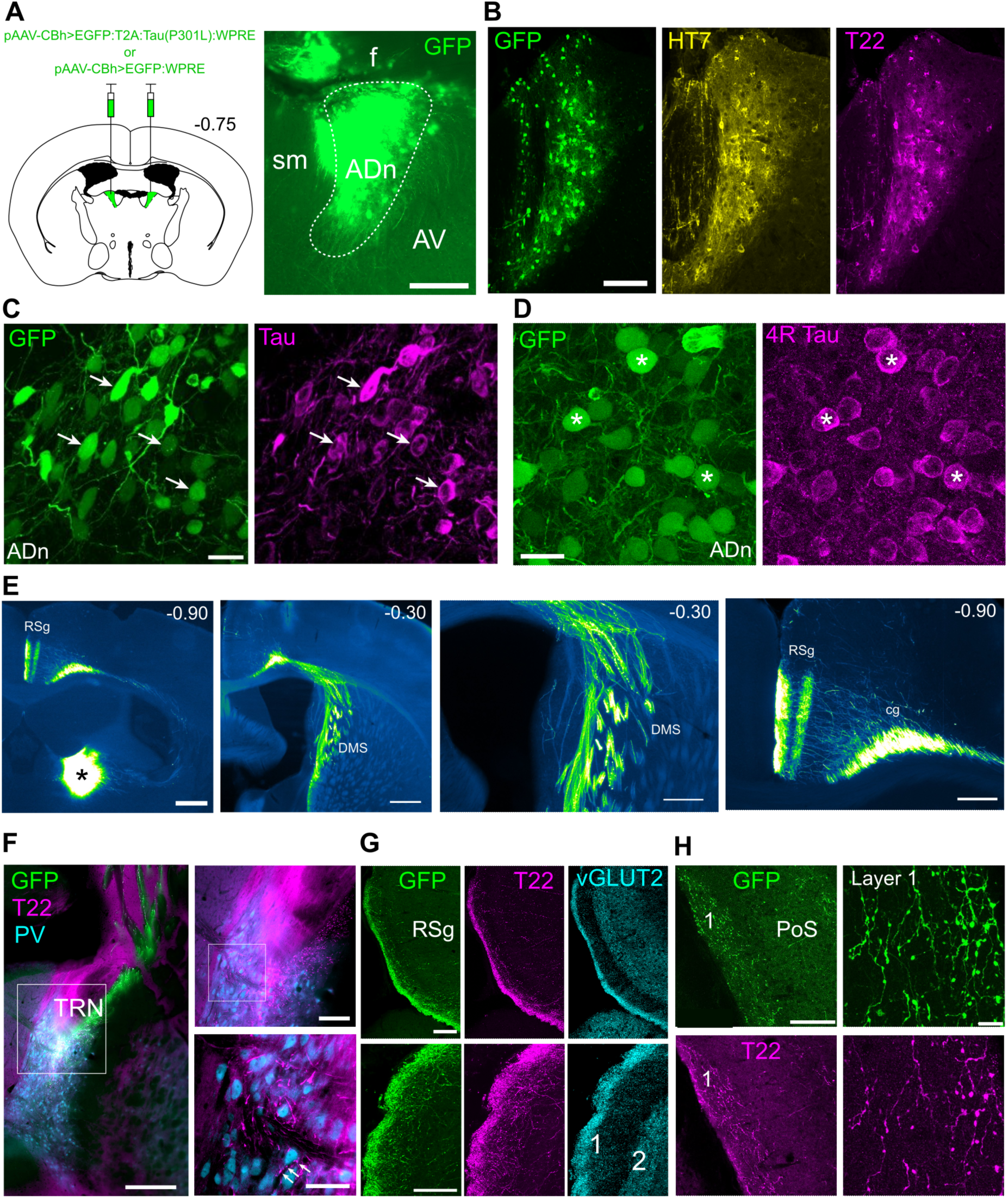
Defining HD-tau mice by htau expression in the HD network. (**A**) Left, schematic of bilateral AAV injections into the ADn (coronal section) at ∼-0.75 mm from Bregma. Right, widefield epifluorescence image of a coronal brain section at the injection site showing GFP expression (green) within the ADn. Case SJ22. (**B**) Viral expression of GFP (green) and human tau, detected by HT7 (yellow) and T22 (oligomeric tau, magenta). Case SH38, 21 µm thick confocal z-projection. (**C**) Colocalization of GFP (green) and phospho-Tau (T231, magenta) in the ADn (e.g. arrows). Case TV178, 29 µm thick confocal z-projection. (**D**) Colocalization of GFP and 4R Tau (magenta) in the ADn (e.g. asterisks). Case TV178, 13.4 µm-thick confocal z-projection. (**E**) Trajectory of GFP-expressing ADn cell axons to postsynaptic target regions. Widefield epifluorescence micrographs, case SH30. Left, injection site centered on the ADn (asterisk, over-exposed) at -0.90 mm from Bregma. Axons travel lateral and rostral into the TRN followed by the DMS (-0.30 mm from Bregma), then cross the corpus callosum, entering the cingulum bundle traveling caudally to branch in the RSg (far right, -0.90 mm from Bregma). (**F**) Left, projections from GFP-expressing ADn cells to the TRN followed by the DMS. Right, innervation of the TRN by GFP+ T22+ axons. Inset top right, detail of T22+ axons in the dorsal TRN, which contains PV-immunoreactive neurons (cyan). Bottom right inset, detail of innervated TRN cells (e.g. arrows). Case SJ14, widefield epifluorescence. (**G**) Top, innervation of the RSg by GFP-expressing axons from the ADn, with GFP (green) and T22 (magenta) localized to the vGLUT2-immunoreactive plexus (cyan) in layer 1. Case SH38, 28 µm thick confocal maximum intensity z-projection. Bottom, detail of ADn terminals in layer 1 of RSg, 25.5-µm thick confocal z-projection. (**H**) Left, innervation of the PoS by GFP+ T22+ axons from the ADn. Case SH38, 20.6 µm thick confocal z-projection. Right, detail of ADn terminals in layer 1 of PoS, 25.9 µm thick confocal maximum intensity z-projection. Scale bars (µm): A 100; B 100; C 20; D 20; E left two panels 500, right two panels 250; F 300, 100, 50; G 100; H left 100, right 10. Abbreviations: sm, stria medullaris; f, fimbria. See also Fig. S1.

**Table 1.**
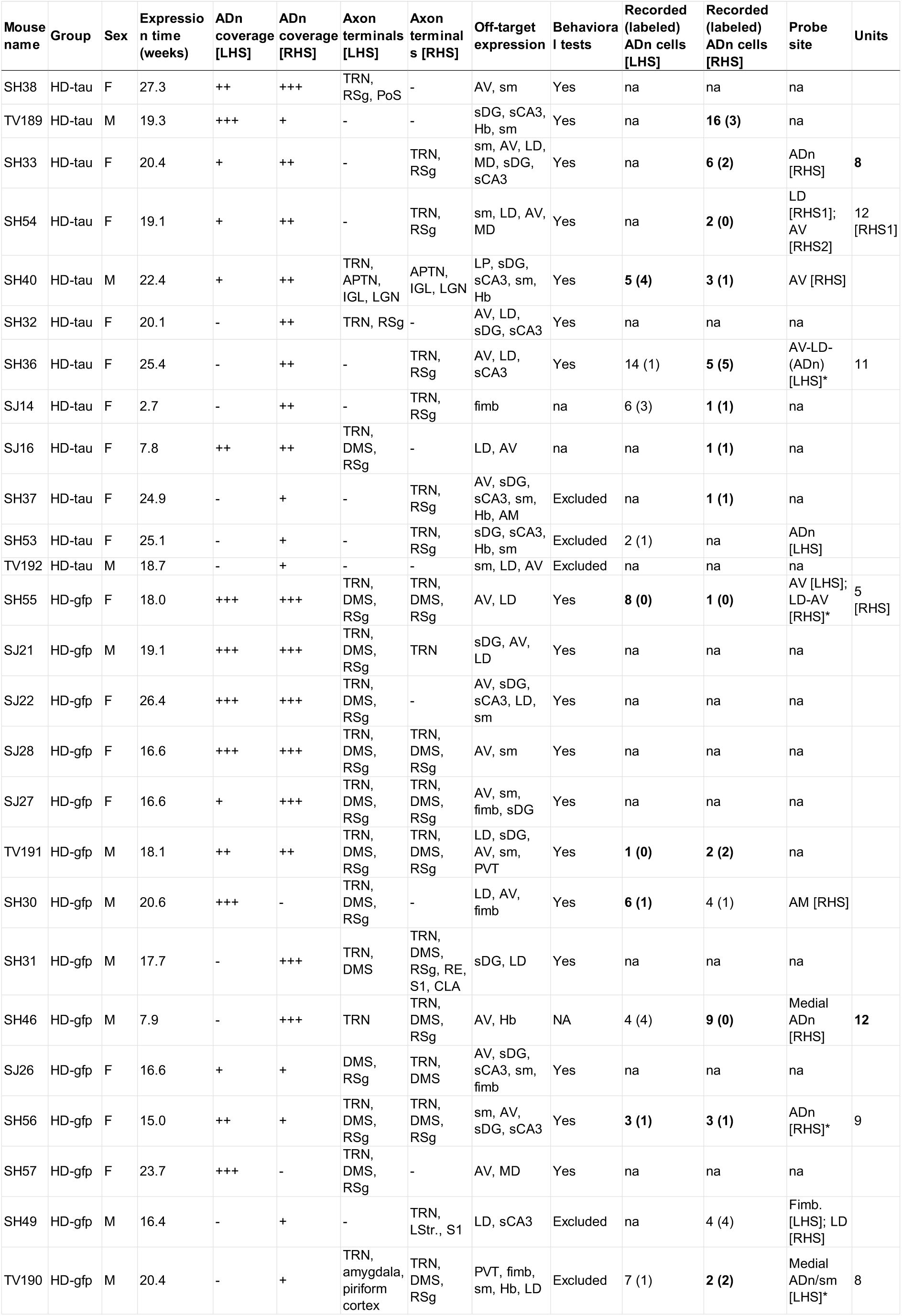
HD-tau and HD-gfp mice included for behavioral testing and/or *in vivo* recordings. Expression time refers to time between virus injection and transcardial perfusion; *in vivo* recordings were performed 1-3 days before perfusion. ADn coverage refers to viral expression of GFP in somata throughout the ADn based on the following scores: – lack of expression; + sparse expression; ++ moderate expression; +++ abundant expression. Axon terminals were observed in the listed target regions but not all regions were examined in each mouse; some mice had low levels of GFP expression in axons. Recorded cell and unit counts are shown in bold if they were from hemispheres with GFP expression within the ADn. Abbreviations: F, female; M, male; LHS, left hemisphere; RHS, right hemisphere; na, not applicable/available; ADn, anterodorsal thalamic nucleus; TRN, thalamic reticular nucleus; DMS, dorsomedial striatum; RSg, granular retrosplenial cortex; PoS, postsubiculum; AV, anteroventral thalamus; LD, laterodorsal thalamus; sm, stria medullaris; sDG, septal pole of the dentate gyrus; sCA3, septal pole of the CA3 region of the hippocampus; Hb, habenula; MD, mediodorsal thalamus; LP, lateroposterior thalamus; AM, anteromedial thalamus; PVT, paraventricular thalamus; fimb, fimbria; APTN, anterior pretectal nucleus; IGL, intergeniculate leaflet; LGN, lateral geniculate nucleus; RE, reuniens nuclear complex; S1, primary somatosensory cortex; CLA, claustrum; LStr., lateral striatum. Asterisks: lack of GFP in probe recording location.

| Mouse name | Group | Sex | Expression time (weeks) | ADn coverage [LHS] | ADn coverage [RHS] | Axon terminals [LHS] | Axon terminals [RHS] | Off-target expression | Behavioral tests | Recorded (labeled) ADn cells [LHS] | Recorded (labeled) ADn cells [RHS] | Probe site | Units |
| --- | --- | --- | --- | --- | --- | --- | --- | --- | --- | --- | --- | --- | --- |
| SH38 | HD-tau | F | 27.3 | ++ | +++ | TRN, RSg, PoS | - | AV, sm | Yes | na | na | na |  |
| TV189 | HD-tau | M | 19.3 | +++ | + | - | - | sDG, sCA3, Hb, sm | Yes | na | **16 (3)** | na |  |
| SH33 | HD-tau | F | 20.4 | + | ++ | - | TRN, RSg | sm, AV, LD, MD, sDG, sCA3 | Yes | na | **6 (2)** | ADn [RHS] | **8** |
| SH54 | HD-tau | F | 19.1 | + | ++ | - | TRN, RSg | sm, LD, AV, MD | Yes | na | **2 (0)** | LD [RHS1]; AV [RHS2] | 12 [RHS1] |
| SH40 | HD-tau | M | 22.4 | + | ++ | TRN, APTN, IGL, LGN | APTN, IGL, LGN | LP, sDG, sCA3, sm, Hb | Yes | **5 (4)** | **3 (1)** | AV [RHS] |  |
| SH32 | HD-tau | F | 20.1 | - | ++ | TRN, RSg | - | AV, LD, sDG, sCA3 | Yes | na | na | na |  |
| SH36 | HD-tau | F | 25.4 | - | ++ | - | TRN, RSg | AV, LD, sCA3 | Yes | 14 (1) | **5 (5)** | AV-LD- (ADn) [LHS]* | 11 |
| SJ14 | HD-tau | F | 2.7 | - | ++ | - | TRN, RSg | fimb | na | 6 (3) | **1 (1)** | na |  |
| SJ16 | HD-tau | F | 7.8 | ++ | ++ | TRN, DMS, RSg | - | LD, AV | na | na | **1 (1)** | na |  |
| SH37 | HD-tau | F | 24.9 | - | + | - | TRN, RSg | AV, sDG, sCA3, sm, Hb, AM | Excluded | na | **1 (1)** | na |  |
| SH53 | HD-tau | F | 25.1 | - | + | - | TRN, RSg | sDG, sCA3, Hb, sm | Excluded | 2 (1) | na | ADn [LHS] |  |
| TV192 | HD-tau | M | 18.7 | - | + | - | - | sm, LD, AV | Excluded | na | na | na |  |
| SH55 | HD-gfp | F | 18.0 | +++ | +++ | TRN, DMS, RSg | TRN, DMS, RSg | AV, LD | Yes | **8 (0)** | **1 (0)** | AV [LHS]; LD-AV [RHS]* | 5 [RHS] |
| SJ21 | HD-gfp | M | 19.1 | +++ | +++ | TRN, DMS, RSg | TRN | sDG, AV, LD | Yes | na | na | na |  |
| SJ22 | HD-gfp | F | 26.4 | +++ | +++ | TRN, DMS, RSg | - | AV, sDG, sCA3, LD, sm | Yes | na | na | na |  |
| SJ28 | HD-gfp | F | 16.6 | +++ | +++ | TRN, DMS, RSg | TRN, DMS, RSg | AV, sm | Yes | na | na | na |  |
| SJ27 | HD-gfp | F | 16.6 | + | +++ | TRN, DMS, RSg | TRN, DMS, RSg | AV, sm, fimb, sDG | Yes | na | na | na |  |
| TV191 | HD-gfp | M | 18.1 | ++ | ++ | TRN, DMS, RSg | TRN, DMS, RSg | LD, sDG, AV, sm, PVT | Yes | **1 (0)** | **2 (2)** | na |  |
| SH30 | HD-gfp | M | 20.6 | +++ | - | TRN, DMS, RSg | - | LD, AV, fimb | Yes | **6 (1)** | 4 (1) | AM [RHS] |  |
| SH31 | HD-gfp | M | 17.7 | - | +++ | TRN, DMS | TRN, DMS, RSg, RE, S1, CLA | sDG, LD | Yes | na | na | na |  |
| SH46 | HD-gfp | M | 7.9 | - | +++ | TRN | TRN, DMS, RSg | AV, Hb | NA | 4 (4) | **9 (0)** | Medial ADn [RHS] | **12** |
| SJ26 | HD-gfp | F | 16.6 | + | + | DMS, RSg | TRN, DMS | AV, sDG, sCA3, sm, fimb | Yes | na | na | na |  |
| SH56 | HD-gfp | F | 15.0 | ++ | + | TRN, DMS, RSg | TRN, DMS, RSg | sm, AV, sDG, sCA3 | Yes | **3 (1)** | **3 (1)** | ADn [RHS]* | 9 |
| SH57 | HD-gfp | F | 23.7 | +++ | - | TRN, DMS, RSg | - | AV, MD | Yes | na | na | na |  |
| SH49 | HD-gfp | M | 16.4 | - | + | - | TRN, LStr., S1 | LD, sCA3 | Excluded | na | 4 (4) | Fimb. [LHS]; LD [RHS] |  |
| TV190 | HD-gfp | M | 20.4 | - | + | TRN, amygdala, piriform cortex | TRN, DMS, RSg | PVT, fimb, sm, Hb, LD | Excluded | 7 (1) | **2 (2)** | Medial ADn/sm [LHS]* | 8 |

The ADn was localized by high levels of lipofuscin and biotin (Fig. S1B), which correlated with C1ql2 immunoreactivity (Fig. S1A) (Roy et al., 2021; Kapustina et al., 2024). We detected htau in ADn somata and dendrites in HD-tau mice but not in HD-gfp mice based on immunoreactivity to HT7 (human-specific tau), T22 (oligomeric tau), 4R tau, and phospho-tau T231 (Fig. 1B-D, S1D). The oligomeric, pre-aggregated form is considered to be a driver of early pathological changes (Lasagna-Reeves et al., 2011; Lasagna-Reeves et al., 2012; Taddei et al., 2023). Approximately 90% of GFP+ cells in the ADn showed positive immunoreactivity for T22 in HD-tau mice (96 tested cells from 4 mice; Fig. 1B, S1E) while no T22+ cells were detected in HD-gfp mice (Fig. S1D).

Axons and presynaptic terminals tested in the TRN, RSg, and PoS were immunoreactive for oligomeric tau in HD-tau mice but not in HD-gfp mice (Fig. 1F-H, S1F-G). The htau+ axons in postsynaptic target regions were GFP+ (case SH38; Fig. 1G-H). We did not detect htau+ cells lacking GFP in postsynaptic regions of the ADn, suggesting a lack of transsynaptic spread of htau (Dujardin et al., 2022). In the TRN, ADn axons formed basket-like terminals around parvalbumin (PV)-expressing neurons (Fig. 1F). In the RSg, ADn axon terminals formed a plexus in the most superficial part of layer 1 and colocalized with vesicular glutamate transporter 2 (vGLUT2; Fig. 1G). A similar pattern was observed in the PoS (Fig. 1H). These innervation patterns are consistent with the pattern of ptau immunoreactivity detected in human ADn, TRN, and RSg at early Braak stages (Sárkány et al., 2024). Taken together, these data demonstrate that HD-tau mice express htau within the HD network.

### Disorientation-related alterations to spatial memory in HD-tau mice

Spatial memory is dependent on the HD network (Warburton et al., 1997; Warburton et al., 2001; Vann, 2013; Frost et al., 2021). To establish whether expression of mutant htau in the HD network affects spatial navigation and orientation, we trained HD-tau and HD-gfp mice in spatial-reference and spatial-working memory tasks after 9-13 weeks of viral expression (Fig. 2, 3, S2).

**Figure 2.**
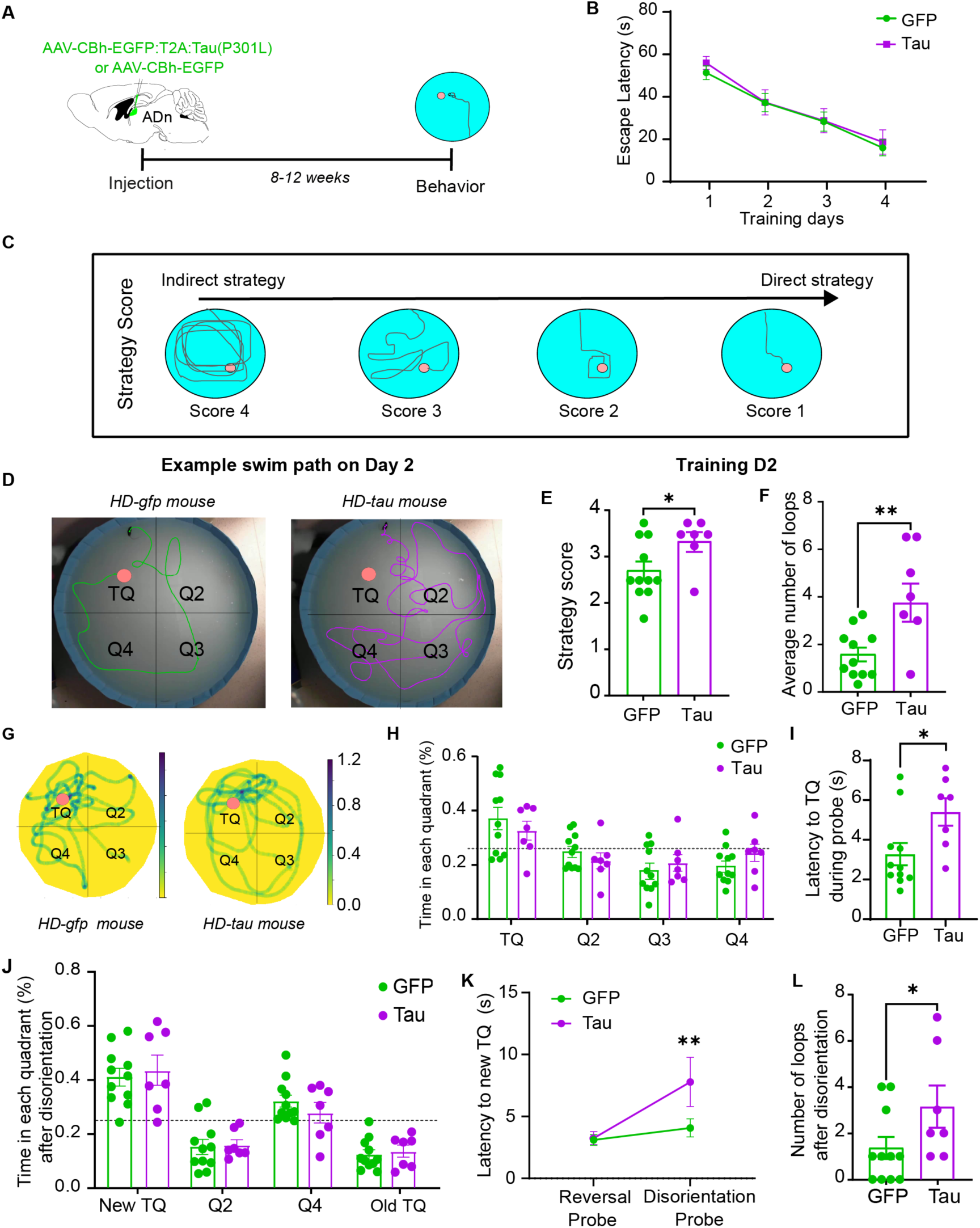
HD-tau mice exhibit disorientation during a spatial reference memory task. (**A**) Experimental design: AAVs were bilaterally injected into the mouse ADn. Following an expression time of 8-12 weeks, behavioral testing was initiated. (**B**) Spatial reference memory was assessed in a classical MWM by measuring the latency to find a hidden platform with a fixed location on 4 consecutive training days (T1-4). (**C**) Schematic of the different swim strategies used to find the hidden platform (Score 4, most indirect target strategy; score 1, most direct target strategy; see Methods). (**D**) Example images of swim paths for an HD-gfp mouse (left) and an HD-tau mouse (right) on training day 2. Note the greater number of loops performed by the HD-tau mouse. (**E**) HD-tau mice showed a significantly lower strategy score (average score across the 4 trials) on training day 2 compared to HD-gfp mice. (**F**) HD-tau mice showed an increase in the number of loops during training day 2. (**G**) Example heatmaps of the swimming activity during the probe test. Both groups of mice spent a significantly longer time in the quadrant normally containing the hidden platform. Scale bar: occupancy in seconds. (**H**) Quantification of the time spent in each quadrant, showing no difference between groups. Both groups spent more time in the TQ compared to chance level. (**I**) HD-tau mice exhibited an increased latency to reach the TQ during the probe test. (**J**) There was no difference observed in the time spent in the new TQ in the probe test after disorientation. Dashed line, chance level. (**K**) Latency to the new TQ in the disorientation probe test compared to in the reversal probe test. (**L**) HD-tau mice exhibited increased looping behavior after disorientation. See also Fig. S2 and Videos S1-S7.

**Figure 3.**
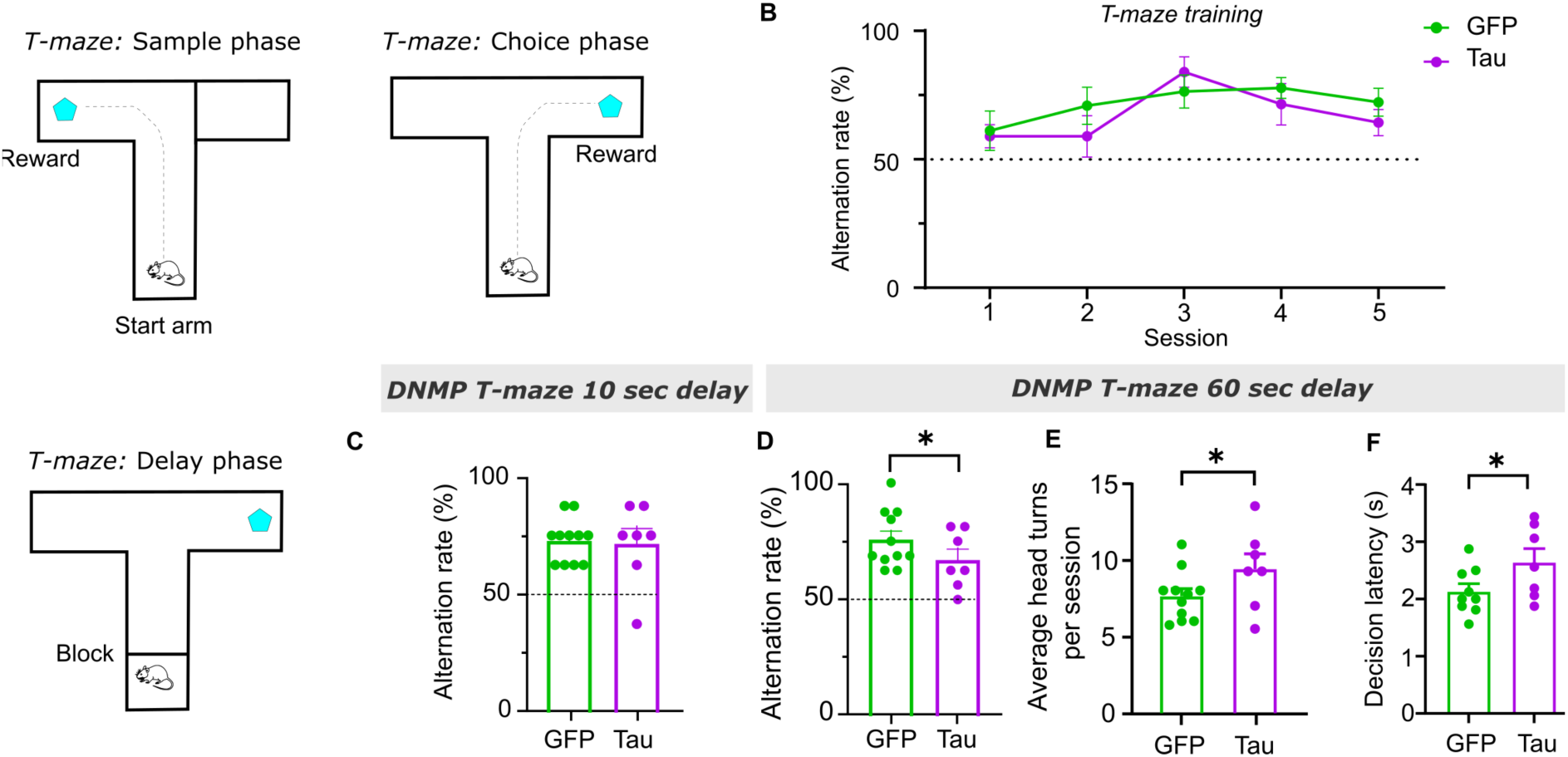
HD-tau mice exhibit features of disorientation during a spatial working memory task. (**A**) Schematic of the non-match-to-place T-maze task. Training consisted of 5 sessions (defined as a block of 8 trials over 2 days) conducted over 10 days. Each trial included 1 sample phase (with one arm blocked and the other containing a water reward), and 1 choice phase (both arms open and only the previously unvisited arm now rewarded). At the end of the training period, mice had a delay interval interposed following the sample phase (delayed non-match-to-place (DNMP) T-maze), which included one session with a 10 s delay (8 trials) followed by two sessions with a 60 s delay (16 trials). (**B**) Both groups learned the task, as shown by the increased alternation rate over time. (**C**) The two groups had similar alternation rates for the 10 s DNMP task. (**D**) There was a significant impairment in the performance of HD-tau mice compared to HD-gfp mice for the 60 s DNMP task (one-tailed t-test). (**E-F**) Quantification of the average number of head turns per session (E) and decision latency (F) for the 60 s DNMP task, showing an increase in both head turns and latency to make a decision for HD-tau mice. Dashed lines, chance level.

Spatial reference memory was assessed using the Morris Water Maze (MWM), whereby mice had to learn to find a hidden platform using spatial cues (Fig. 2B-F, S2B-D). This was followed by a probe test whereby the platform was removed and mice explored the MWM for 60 s (Fig. 2G-I, S2E-G). Both groups learned to find the platform during training, suggesting that HD-tau mice have unimpaired spatial learning with respect to fixed distal visual cues (*training effect (main effect of day)* two-way repeated measures ANOVA: n = 7 HD-tau mice and n = 11 HD-gfp mice, *F*_3,48_ = 37.57, \*\*\**p* < 0.0001; Fig. 2B).

To assess spatial learning during training, we ranked their average search strategies from ‘indirect’ to ‘direct’, based on a scoring system (Berkowitz et al., 2018; Vantomme et al., 2020) (Fig. 2C). On the second day of training, HD-gfp and HD-tau mice favored different strategies, with HD-tau mice using more indirect strategies (*t*_(16)_ = 2.207, \**p* = 0.042; Fig. 2D). One major difference we observed was that HD-tau mice tended to make several localized loops, mostly in the same direction (clockwise or counterclockwise), while searching for the platform (Fig. 2E, S2B; Videos S1-5). The HD-tau mice made a significantly greater number of loops across four trials compared to HD-gfp mice (median [inter-quartile range, IQR]: 4 [3-6.5] loops for HD-tau mice versus 1.25 [0.75-2.25] loops for HD-gfp mice; *U* = 10.5, \*\**p* = 0.0093; Fig. 2F; Videos S1-5). The average number of loops positively correlated with htau/GFP expression levels in HD-tau mice but not in HD-gfp mice (Spearman *r* = 0.821 for HD-tau mice versus *r* = 0.087 for HD-gfp mice; Fig. S2C). No differences were detected in their swimming speeds (Fig. S2D). The increased looping behavior likely contributed to the less direct strategies used by HD-tau mice. This difference was resolved by day 3, suggesting HD-tau mice display a difference in the initial acquisition of the spatial map when compared to HD-gfp mice (Fig. S2B).

In the probe test, both groups spent more time in the target quadrant (TQ), which contains the hidden platform, compared to chance, with no differences detected between groups (HD-gfp vs chance*: U =* 7, \*\*\**p* = 0.0002; HD-tau vs chance: *U* = 7, \**p* = 0.0169, HD-gfp vs HD-tau: *t*_(16)_ = 0.7705, *p* = 0.4522; Fig. 2G-H), indicating that both groups of mice learned the location of the platform. However, HD-tau mice took a significantly longer time to reach the TQ (mean ± SEM: 5.4 ± 0.7 s for HD-tau mice versus 3.3 ± 0.6 s for HD-gfp mice, *t*_(16)_ = 2.373, *\*p* = 0.0305; Fig. 2I). No differences were detected for the number of entries (Fig. S2E), total distance traveled (Fig. S2F), or swim speed (Fig. S2G) in the TQ. Next, we tested the mice in a reversal learning protocol, whereby the platform location was swapped to the opposite quadrant of the pool (new TQ). We detected no differences in the performance between groups for both the training trials and post-reversal probe test (Fig. S2H-L).

To test whether the increased looping behavior in HD-tau mice was linked to alterations in spatial orientation, we tested the same mice again in the MWW immediately after disorienting them. This was achieved by placing each mouse in a small opaque box, rotating the box 16 times, then immediately placing them in the MWM (Fig. S2A, Methods). Although both groups spent a similar proportion of time in the new TQ, remaining above chance levels (HD-gfp vs chance*: U* = 6, \*\**p* = 0.0040; HD-tau vs chance: *U* = 6, \**p* = 0.0239, HD-gfp vs HD-tau: *t*_(16)_ = 0.4040, *p* = 0.6916; Fig.2K), the disorientation procedure reduced the time spent in the new TQ (Fig. S2M-N). Interestingly, in the disorientation probe test, HD-tau mice now required significantly more time to reach the new TQ compared to their performance in the reversal probe test (*group × timepoint* two-way repeated measures ANOVA: *F*_1,16_ = 5.469, *\*p* = 0.0327, *post-hoc* LSD test for HD-tau group: *\*\*p* = 0.0019; Fig. 2K); this was not observed for HD-gfp mice (*p* = 0.3765; Fig.2K). The HD-tau mice also exhibited more looping behavior compared to HD-gfp mice following disorientation (*U* = 18.50, \**p* = 0.0326; Fig. 2L; Videos S6-7).

Finally, we performed a long-term spatial reference memory test by carrying out another probe test in the MWM one month after the disorientation probe test (Fig. S2A) and found that both groups spent significantly longer time in the new TQ (Fig. S2O), suggesting that HD-tau mice have unimpaired retrieval of long-term spatial memories.

Spatial working memory was tested using a rewarded alternation non-matching-to-place T-maze task (Fig. 3). Both groups learned the task (two-way repeated measures ANOVA; n = 7 HD-tau mice and n = 11 HD-gfp mice; *training effect F*_4,64_ = 3.342, *\*p* = 0.0151; *training x group effect F*_4,64_ = 0.8354, *p* = 0.5077; Fig. 3B). When we introduced an additional 10 s delay between sample and choice runs, both groups again performed similarly (*t*_(16)_ = 0.2081, *p* = 0.8378; Fig. 3C). However, when we increased the difficulty by introducing a 60 s delay, the alternation rate of HD-tau mice was significantly reduced compared to HD-gfp mice (66.9 ± 4.6% for HD-tau mice vs 75.7 ± 3.7% for HD-gfp mice, *t*_(16)_ = 1.887, *\*p* = 0.0387; Fig. 3D), consistent with previous findings of selective silencing of the ADn (Roy et al., 2021; Roy et al., 2022).

Interestingly, during the choice phase of the T-maze task with a 60 s delay, HD-tau mice made more head turns than HD-gfp mice in the start arm before reaching the decision point (2.3 ± 0.3 head turns for HD-tau mice vs 1.8 ± 0.1 head turns for HD-gfp mice, *t*_(12)_ = 1.970, *\*p* = 0.0362; Fig. 3E), and took significantly longer to make a decision (2.7 ± 0.3 s for HD-tau mice vs 2.2 ± 0.1 s for HD-gfp mice, *t*_(12)_ = 1.879, *\*p* = 0.0424; Fig. 3F). These behavioral patterns may be related to spatial disorientation.

Mice were also tested for their short-term spatial memory in a 1-trial Y-maze spatial novelty preference task (Sanderson et al., 2007). Both HD-tau and HD-gfp mice showed an equivalent preference for the novel arm and no difference was detected between groups (Fig. S2P). We did not detect any differences between HD-tau and HD-gfp mice in terms of locomotor activity or anxiety-related behavior (Fig. S2Q). Taken together, the behavioral tests revealed that HD-tau mice had subtle alternations to spatial memory and some behavior indicative of disorientation.

### Reduced directionality of ADn cells in HD-tau mice

We hypothesized that the alterations to spatial memory acquisition and recall in HD-tau mice were associated with htau affecting the firing patterns of ADn cells. To test this, we targeted the ADn with glass electrodes in the previously behaviorally-tested mice. To detect HD cells, we passively rotated head-restrained mice on a turntable with respect to fixed visual cues, which provides rotational vestibular and visual inputs required for HD cells (Fig. 4A and S3A) (Blanco-Hernández et al., 2024). Following single-neuron extracellular recordings, we juxtacellularly labeled selected cells with neurobiotin. *Post-hoc* recovery of the labeled cells enabled us to determine the relative recording locations of unlabeled cells, thus providing a robust classification of ADn cells with respect to GFP expression and htau immunoreactivity (Table 1; Fig. S3A). Juxtacellular labeling preserved the expression of htau and GFP (Fig. 4B).

**Figure 4.**
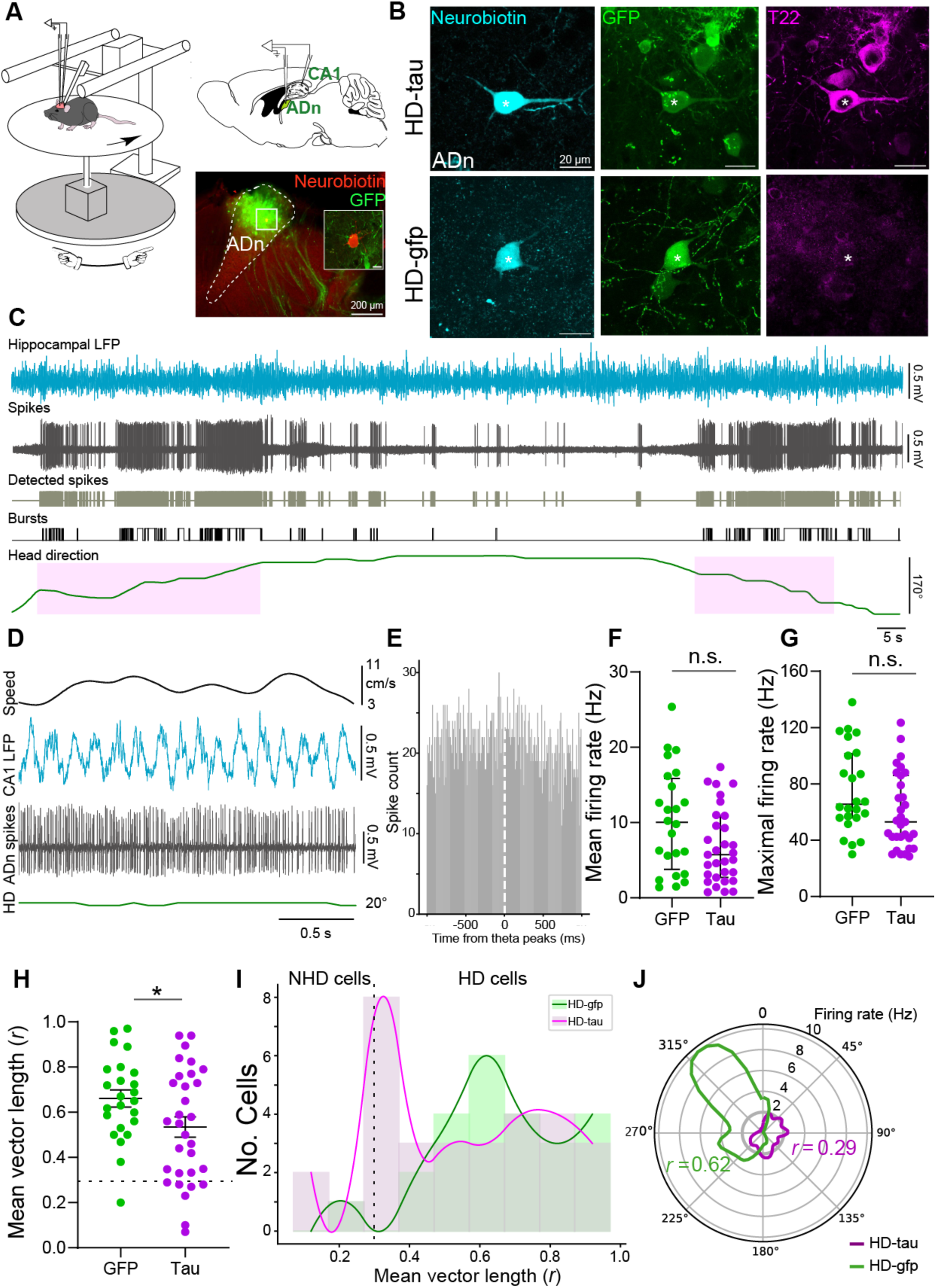
Reduced directionality of ADn cells from HD-tau mice. (**A**) Left, schematic of *in vivo* extracellular recordings in head-restrained mice. The turntable (gray) is passively rotated. The mouse is able to spontaneously run and rest on a running disc. Top right, sagittal brain section indicating target locations of glass electrodes. Bottom right, epifluorescence micrograph of a coronal brain section showing viral expression of GFP (green) in the ADn. Inset, a neurobiotin-labeled HD cell (red, cell SJ14-10), confocal maximum intensity z-projection (12.6 µm thick), scale bar 10 µm. (**B**) Top, recovery of a juxtacellularly labeled ADn cell (neurobiotin, cyan, SH40-072, asterisk) that expressed GFP (green) and was immunoreactive for oligomeric tau (T22, magenta); confocal maximum intensity z-projection (5.1 µm thick), case SH40. Bottom, a juxtacellularly labeled ADn cell (cell SH49-023, asterisk) that expressed GFP but lacked detectable immunoreactivity for T22; confocal maximum intensity z-projection (11.7 µm thick), case SH49. Scale bars, 20 μm. (**C**) Example of simultaneous recording of the CA1 LFP (cyan), action potentials of an ADn HD cell (SH30-05, magenta, pink), detected bursts (black), and head direction (green). The shaded purple area marks the preferred firing direction. (**D**) Spiking activity of an ADn HD cell (SH33-1-h1) in the preferred firing direction during locomotion with theta oscillations in CA1. The cell lacked theta-rhythmic firing. (**E**) Cross-correlogram between CA1 theta and ADn HD cell spikes (SH33-1-h1), *p* < 0.05. (**F-G**) Electrophysiological properties of ADn cells recorded with glass electrodes (F, mean firing rate; G, maximal firing rate). Not significant, n.s. (**H**) Mean vector length (*r*) for all recorded ADn cells with glass electrodes from HD-tau mice and HD-gfp mice. (**I**) Distribution of mean vector lengths for all recorded ADn cells from HD-tau mice and HD-gfp mice (bin size, *r* = 0.1). (**J**) Polar plot of tuning curves for two representative ADn cells from HD-tau mice (SH40-072, magenta) and HD-gfp mice (TV190-2-i, green). Dashed line, threshold for non-HD cells (NHD) with *r* < 0.3. See also Fig. S3 and S4.

We learned to recognize the ADn by the presence of HD cells that exhibited irregular burst firing and a lack of coupling to hippocampal theta oscillations (Figure 4C-E) (Taube, 1995). We localized 116 single extracellularly recorded cells to the ADn based on the locations of 40 recovered juxtacellularly labeled cells from 16 mice (Table 1; Fig. S3A). We excluded 61 ADn cells due to a lack of GFP/htau expression in the recording locations, resulting in n = 40 and n = 35 ADn cells from HD-tau and HD-gfp mice, respectively. Analysis of all glass-electrode recorded ADn cells revealed a trend for lower mean firing rates of ADn cells from HD-tau mice (5.7 [2.7-10.9] Hz for n = 31 cells from HD-tau mice versus 10.0 [3.7-15.8] Hz for n = 24 cells from HD-gfp mice; Mann-Whitney test, *U* = 266, *p* = 0.0732; Fig. 4F). A similar pattern emerged for the maximal firing rates (53 [41.5-86] Hz from HD-tau mice versus 65.5 [56-101.5] Hz from HD-gfp mice; *U* = 259, *p* = 0.054; Fig. 4G).

When we compared the strength of the directional signal (also known as directionality, indicated by the mean vector length *r*), ADn cells from HD-tau mice exhibited a lower mean directionality (*r* = 0.53 ± 0.04 for n = 31 cells from HD-tau mice versus *r* = 0.66 ± 0.04 for n = 24 cells from HD-gfp mice; *t*_(53)_ = 2.079, \**p* = 0.0425; Fig. 4H). We used a threshold of *r* = 0.3 to define ADn HD cells and ADn non-HD (NHD) cells based on published data from the mouse ATN (Blanco-Hernández et al., 2024). The distribution of mean vector length of HD cells from HD-tau mice shifted toward lower values than those from HD-gfp mice (Fig. 4I), suggesting a significantly lower proportion of ADn HD cells in HD-tau versus HD-gfp mice (46.5% versus 95.8%; Fisher’s exact test, \*\**p* = 0.0072). Two representative tuning curves taken from each group highlight key differences in directionality and mean firing rate (Fig. 4J). These results suggest that ADn cells from HD-tau mice have reduced directional coding.

Next, we analyzed the subset of ADn cells classified as HD cells (*r* > 0.3) and found that htau did not significantly alter their directionality (*r* = 0.61 ± 0.04 for n = 20 cells from HD-tau mice versus 0.68 ± 0.03 for n = 23 cells from HD-gfp mice; *t*_(41)_ = 1.251, *p* = 0.5305; Fig. S3B). The peak and background firing rates were similar (Fig. S3C, S3D), as were the directional tuning widths (103.4 [73.54–225]° from HD-tau mice versus 106.9 [60.8–180]° from HD-gfp mice; *U* = 215, *p* = 0.72; Fig. S3E). We also determined the distribution of preferred firing directions, and HD cells from both groups were similar (Fig. S3F).

To further explore these effects at the population level, we also targeted the ADn with acute silicon probes following successful juxtacellular modulation with a glass electrode (Fig. S3G,H). Consistent with the single-cell recordings, we observed lower directionality of ADn cells within htau-enriched zones compared to control GFP-enriched zones (*r* = 0.32 ± 0.07 for n = 8 units from one HD-tau mouse versus *r* = 0.64 ± 0.05 for n = 9 units from one HD-gfp mouse; *t*_(15)_ = 3.957, \*\**p* = 0.0013; Fig. S3I, Table 1). Based on HD cell selection criterion of population recording data (see Methods), we also found a lower proportion of ADn HD cells from HD-tau mice compared to HD-gfp mice (37.5% versus 88.89%; Fisher’s exact test, \**p* = 0.0498).

### Altered burst firing of HD cells in HD-tau mice

Next, we tested whether there were any differences in firing within the receptive field of HD cells, i.e. when the mouse faces the preferred firing direction of a given HD cell (Fig.5). We observed that HD cells from HD-tau mice had a lower percentage of spikes per burst (n = 25 cells from HD-tau mice versus n = 26 cells from HD-gfp mice; *t*_(49)_ = 2.378, \**p* = 0.0214; Fig 5C,D), an increased duration between bursts (*U* = 198, \**p* = 0.0161; Fig.5H) and a longer average burst cycle period compared to ADn HD cells from HD-gfp mice (*U* = 186, \*\**p* = 0.0082; Fig.5I). This is consistent with the altered bursting patterns of HD cells following conditions that promote disorientation via prolonged rotation of mice (Grieves et al., 2022), suggesting that HD-tau mice exhibit disorientation.

**Figure 5.**
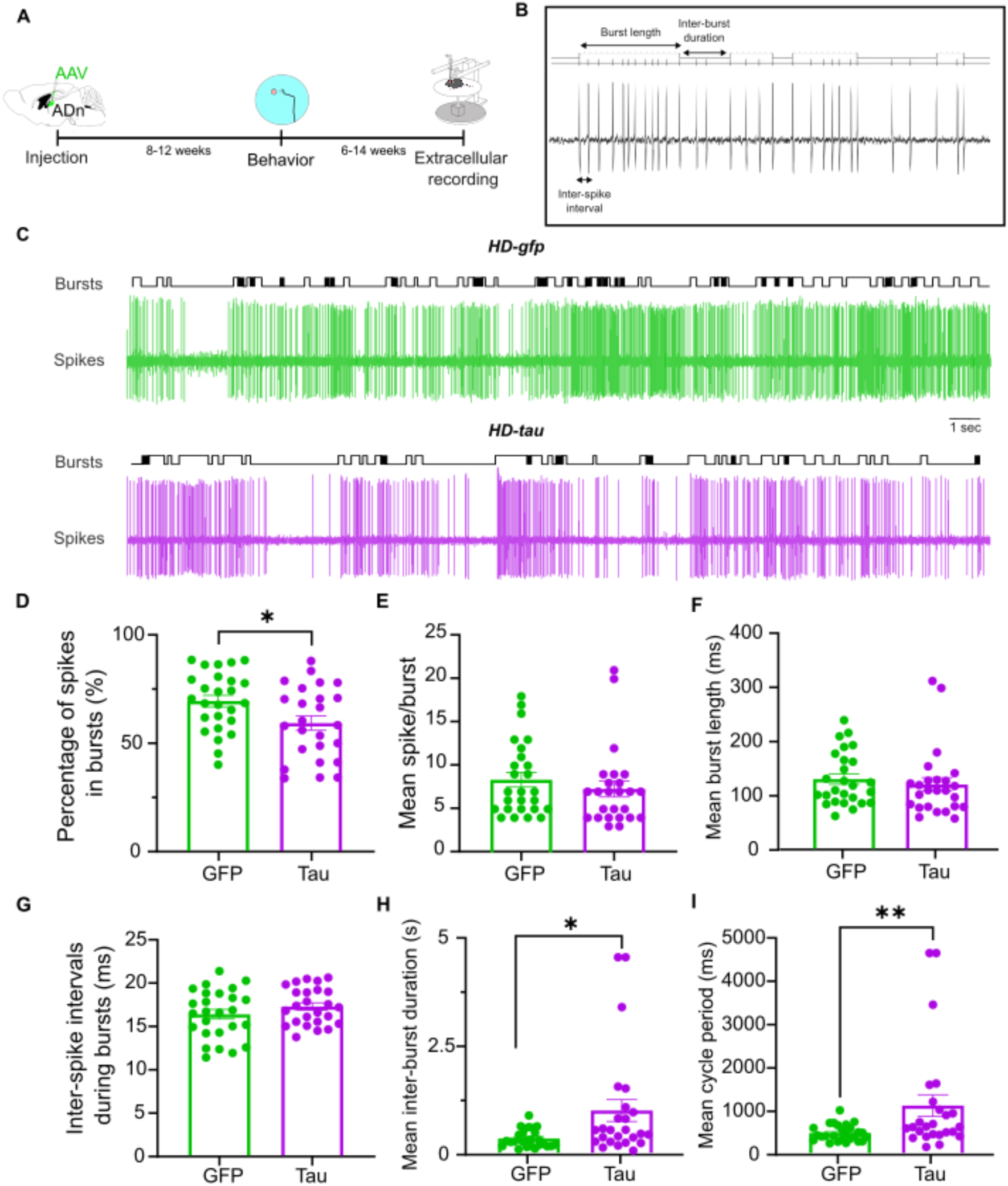
Altered burst firing of HD cells from HD-tau mice. (**A**) Experimental design. Extracellular recordings were performed 6-14 weeks after behavioral testing. The burst analysis was restricted to periods where the mouse was resting and facing the preferred firing direction. (**B**) Example of the different parameters extracted for the burst analysis. (**C**) Example traces of an HD cell from an HD-gfp mouse (top) and an HD cell from an HD-tau mouse (bottom) showing bursting firing patterns. (**D**) HD cells from HD-tau mice had a lower percentage of spikes during bursts (Student’s t-test: n = 25 cells from HD-tau mice versus n = 26 cells from HD-gfp mice; *t*_(49)_ = 2.378, \**p* = 0.0214). (**E-G**) The mean number of spikes per burst (E), as well as the mean duration of bursts (F) and inter-spike interval during bursts were not different across groups, (E) *t*_(49)_ = 0.8634, *p* = 0.3921; (F) *t*_(49)_ = 1.283, *p* = 0.2054; (G) Mann-Whitney test: *U* = 264, *p* = 0.2547. (**H**) The mean duration between bursts was increased in HD-tau mice (*U* = 198, \**p* = 0.0161). (**I**) The average burst cycle period was significantly longer in HD-tau mice compared to HD-gfp mice (*U* = 186, \*\**p* = 0.0082). See also Fig. S4.

Hippocampal theta oscillations, an indicator of spatial-memory related neuronal coordination, were unaffected by htau in the HD network (Fig. S4). Taken together, these data suggest that htau selectively affects the directionality of ADn cells and the firing mode of HD cells within the preferred firing direction. These disrupted firing properties may explain the alterations to spatial orientation and navigation observed in the behavioral tasks.

## Discussion

Spatial disorientation is emerging as an early cognitive biomarker for dementia (Coughlan et al., 2018), which might be explained by early impairments in the HD network. The human ADn contains a high density of ptau (Braak and Braak, 1991b, a; Xuereb et al., 1991), and is even detected in ADn neurons at ‘pre-Braak’ stage 0 (Sárkány et al., 2024), which may affect the function of HD cells and, in turn, synaptic integration by their postsynaptic target neurons. To this end, we virally expressed mutant human tau in the ADn. This resulted in oligomeric and phosphorylated forms of htau accumulating in somata, dendrites, and axons of ADn cells. Axon terminals containing htau were distributed in postsynaptic target areas of the ADn, including the TRN, DMS, RSg, and PoS. Our data show that while HD-tau mice managed to acquire spatial reference and working memory tasks following training, which are well-known hippocampal-dependent forms of spatial memory (Deacon et al., 2002; Reisel et al., 2002), they used different search strategies and exhibited looping behavior during the initial training phase of the MWM task, and showed hesitations in making choices in the T-maze task, which we attributed to the effect of htau on the HD network. By recording in the ADn, we found that ADn cells from HD-tau mice had significantly reduced directionality affecting the total number of HD cells detected. These HD cells also exhibited altered burst firing. These alterations likely affected the coordination of postsynaptic target neurons, such as cortical HD cells (van der Goes et al., 2024), contributing to spatial disorientation during the tasks.

### Pathological tau contributes to spatial disorientation during memory acquisition and recall

The ADn has received much attention lately and has been characterized as a key brain region for spatial navigation (Weiss and Derdikman, 2018; Aggleton and O’Mara, 2022), but to fully understand its role in spatial memory, it needs to be distinguished from the other anterior thalamic nuclei. The ADn is required for integrating local and distal cues (Stackman et al., 2012). Selective optogenetic inhibition of the ADn impairs spatial working memory performance in a delayed non-matching-to-place T-maze task (Roy et al., 2021; Roy et al., 2022) and disrupts memory retrieval following a contextual fear memory paradigm (Roy et al., 2021; Vetere et al., 2021). Selective inactivation of the ADn in rats prior to MWM training increases the latency to the hidden platform during the probe test without affecting swimming speed (Safari et al., 2020).

How does the accumulation of oligomeric tau affect ADn function and spatial learning? We observed main effects in the MWM on the 2^nd^ day of training, when HD-tau mice used different strategies to reach the hidden platform and exhibited a clear looping behavior that was not observed in HD-gfp mice. These observations suggest an impairment in the early but not late phases of spatial memory acquisition. Interestingly, Donato and colleagues showed that in the initial phase of MWM training, hippocampal neurons undergo structural and network-related plasticity that is required for spatial memory acquisition, and that this plasticity is at its highest on the 2^nd^ day of training (Donato et al., 2013). Another study found that the firing rate of hippocampal place cells as well as the accuracy of spatial information increases dramatically between the 1^st^ and 2^nd^ days of incremental spatial learning (Nakashiba et al., 2008). This suggests that htau alters the communication between brain regions required for spatial navigation in HD-tau mice at this critical time, despite the mice being able to successfully acquire this form of spatial reference memory. A likely source of this interference is the RSg, which receives extensive input from the ADn (Shibata, 1993a; van der Goes et al., 2024) and is required for spatial learning (Vann et al., 2009).

The looping behavior observed in HD-tau mice in the MWM is reminiscent of spatial disorientation, as suggested by other studies (Stackman et al., 2012; Grieves et al., 2022). Mice that are disoriented immediately before testing them in a MWM task, by means of clockwise and counterclockwise rotation for 60 s in a closed box, exhibit looping behavior, and are less likely to use spatial strategies to find the hidden platform, and make more heading errors (Stackman et al., 2012). It is noteworthy that in our experiments, HD-tau mice showed increased looping behavior in the MWM even without disorienting them prior to the MWM. Furthermore, the greater number of head turns and increased latency to make decisions in the T-maze task suggests that HD-tau mice also exhibited some form of disorientation when engaging their spatial working memory. Taken together, our data suggest that htau in the HD network can induce disorientation, altering the way in which spatial memories are acquired.

### Effects of htau on head direction cell activity explains disorientation

Directional information is crucial for spatial navigation (van der Meer et al., 2010; Butler et al., 2017). The ADn is a hub for HD cells (Taube, 1995), and an impairment in HD signaling from the ADn can affect the strategy used by mice during spatial learning in the MWM (Vantomme et al., 2020). It was demonstrated that chemogenetic inhibition of the anterodorsal TRN broadens the directional tuning of ADn HD cells and alters egocentric search strategies in the MWM (Vantomme et al., 2020). Grieves and colleagues showed that disorientation, which is observed as looping behavior, causes a decrease in the directionality of ADn HD cells, as well as a decrease in their peak firing rate and burst index score (Grieves et al., 2022). This strongly suggests that the behavioral disorientation experienced by HD-tau mice is due to the reduced directionality of ADn cells as well as their altered burst firing patterns.

Through what mechanisms might ptau affect the activity of ADn cells? Previous work in animal models has shown that ptau can disrupt neuronal and synaptic function. Some studies showed that hyperphosphorylated tau reduces neuronal excitability, alters neuronal oscillatory patterns, prolongs cortical down states, and decreases firing rates (Yoshiyama et al., 2007; Polydoro et al., 2009; Hoover et al., 2010; Menkes-Caspi et al., 2015; Angulo et al., 2017; Hatch et al., 2017; Busche, 2019; Busche et al., 2019). Pathological tau was reported to depolarize the resting membrane potential and increase the depolarizing voltage deflection or “sag” evoked by hyperpolarization. The increased sag potential has been linked to disruptions in dendritic microtubule-dependent transport of hyperpolarization-activated cyclic nucleotide-gated channels (Crimins et al., 2012). Other studies reported that tau secreted into human cerebrospinal fluid induces neuronal hyperexcitability (Brown et al., 2023) and tau oligomers interfere with action potential waveforms, increasing firing rates (Hill et al., 2019; Hill et al., 2021).

Firing patterns of HD cells, including burst firing, have been suggested to support both the precision and persistence necessary to encode HD information over multiple brain regions. A malformation of the horizontal semicircular canals can induce bursting in the ADn, leading to an unstable HD signal (Valerio and Taube, 2016). Inputs from the ATN, CA1 and dorsal subiculum have been shown to converge on layer 5 RSg pyramidal neurons (Yamawaki et al., 2019). The observed decrease in the average number of spikes per burst and increase in the inter-burst duration for HD cells from HD-tau mice potentially affects the synaptic integration of cortical and thalamic inputs in the RSg, which we predict would affect the timing of spatially-responsive cells such as RSg HD cells, and may also impact HD cell-dependent encoding of allocentric boundary cells and grid cell firing (Burgess et al., 2007; Burgess, 2008; Hasselmo and Brandon, 2008; Alexander et al., 2023). Brennan and colleagues used biophysical models to show that low-rheobase neurons but not regular-spiking neurons in the retrosplenial cortex encode persistent input from afferent PoS HD cells (Brennan et al., 2020). The authors suggested that the specific intrinsic properties of low-rheobase neurons improve their encoding of changing inputs. Simonnet and colleagues found that the bursting activity of neurons in subicular principal cells correlates with encoding of spatial information, revealing that bursts induce sharper firing fields and convey more spatial information than regular spikes (Simonnet and Brecht, 2019). Burst firing has also been found to affect the degree of cortical responsiveness (Fanselow et al., 2001; Swadlow and Gusev, 2001). Taken together, burst firing has been shown to be functionally relevant to spatially tuned neurons, possibly by serving as a mechanism to transmit spatial information to downstream structures.

It is interesting to note that in another study that reported spatial memory deficits following ATN lesions, the authors detected decreased bursting in cells of the subiculum, a known postsynaptic region of the ATN, with a decrease in the average spikes per burst and an increase in the inter-burst interval (Frost et al., 2021). Inactivation or lesions of the ATN were shown to significantly compromise grid cell function in the entorhinal cortex, emphasizing the interdependence of these spatial navigation systems (Winter et al., 2015). Accordingly, the lower directionality of ADn cells in HD-tau mice implies that htau might have downstream effects, leading to impaired firing in postsynaptic target regions.

### Vulnerability of the head direction network

What makes the ADn so vulnerable to tau pathology and neurodegeneration (Braak and Braak, 1991a; Xuereb et al., 1991)? And why specifically the ADn and not the entire ATN? We propose that this is due to a combination of persistently high firing rates, maintenance of large axonal arbors, and high rates of autophagy. First, compared to the cortex, the ADn contains a high density of HD cells, which fire at high rates within the preferred firing direction (Taube, 1995), with little accommodation or adaptation (Taube and Bassett, 2003). We hypothesize that when the head is facing the same direction for a prolonged time period, the subpopulations of HD cells in the ADn for this preferred firing direction would need to maintain high firing rates (Shine et al., 2016; Kim and Maguire, 2019). A change in head position would presumably shift this demand onto other cells. The very high levels of biotin we observe in the mouse ADn, in contrast to the AV (Fig. S1), are consistent with other neurons that maintain high rates (such as ‘fast spiking’ GABAergic neurons) (Salib et al., 2019).

Second, large axonal arbors with hundreds of thousands of presynaptic terminals would place high metabolic demands on a cell. This is the case for the dopaminergic neurons of the substantia nigra, which have enormous axonal arbors and are highly vulnerable to the accumulation of aggregates of alpha-synuclein, leading to neurodegeneration in Parkinson’s disease (Pissadaki and Bolam, 2013). Although no studies have reported full single-neuron reconstructions of ADn cells (to our knowledge), we predict that individual ADn cells have extensive and multiple-branching axons, which is supported by the observation of multiple target regions and the dense plexus of small terminals in the superficial RSg and PoS (Fig. 1, S1). Third, neurons with a high energy demand require high rates of clearance of metabolic waste products via autophagy, with the most vulnerable neurons having the highest axon to soma cytoplasmic volume ratio (Nixon and Rubinsztein, 2024). We observed high levels of lipofuscin in the mouse ADn compared to adjacent regions (Fig. S1), suggesting a demand on the autophagy-lysosomal pathway, which may also include degradation of ptau retrogradely transported from the axon terminals. The AV, which also projects to L1 of the RSg (Sripanidkulchai and Wyss, 1986), has a lower density of HD cells that are distributed mainly in the medial part of AV closest to the ADn (Tsanov et al., 2011; Lomi et al., 2023). The human AV is relatively resistant to neurodegeneration, and accumulates ptau only in late Braak stages (Xuereb et al., 1991; Sárkány et al., 2024). The high metabolic demand of ADn cells may contribute to the observation that the retrosplenial cortex and cingulum bundle are the earliest sites to exhibit glucose hypometabolism in early Alzheimer’s disease (Minoshima et al., 1997; Villain et al., 2008).

### Limitations

The majority of tauopathy animal models exhibit wide-ranging cortical pathology, making it difficult to confidently link any detectable cognitive impairment to a particular brain region or cell type (Viney et al., 2022). To overcome the limitations of transgenic lines, we introduced mutant htau directly into the ADn, a major hub of the HD network. Although this study offers important insights into the role of htau on firing properties of HD cells and spatial memory, it is limited by the accuracy of targeting the ADn and the proportion of labeled ADn cells. This meant excluding mice following behavioral testing and *in vivo* recordings, due to lack of expression in the ADn and/or mis-targeting the silicon probe. Despite our strict inclusion criteria, we cannot rule out that some off-target expression (Table 1) may have contributed to certain behavioral changes in some of the mice. Viral expression did not cover the entirety of the ADn, so we expect effect sizes to be larger by using a more refined genetic-targeting approach. While our data focused on the ADn as a hub within the HD network, other regions, which contain spatially responsive cells such as grid cells or border cells may also be affected by htau expression (Fu et al., 2017). Future studies investigating firing properties of postsynaptic neurons in the RSg, PoS or the entorhinal cortex are required to determine the effects on the entire network.

## Conclusion

We have demonstrated that the accumulation of htau in the HD network induces alterations in the firing patterns of HD cells, which we suggest promotes disorientation, affecting spatial memory performance. Importantly, HD-tau mice still learned the spatial tasks; htau affected the way in which HD-tau mice learned or performed the tasks rather than what they learned. Subtle behavioral changes in spatial navigation and orientation in aged individuals may serve as a compelling early cognitive biomarker for predicting mild cognitive impairment or dementia. Earlier detection of at-risk individuals could enhance opportunities for intervention through dietary modifications or other lifestyle changes, as well as early pharmacological and immunological treatment strategies.

## Supplementary Figures

**Figure S1.**
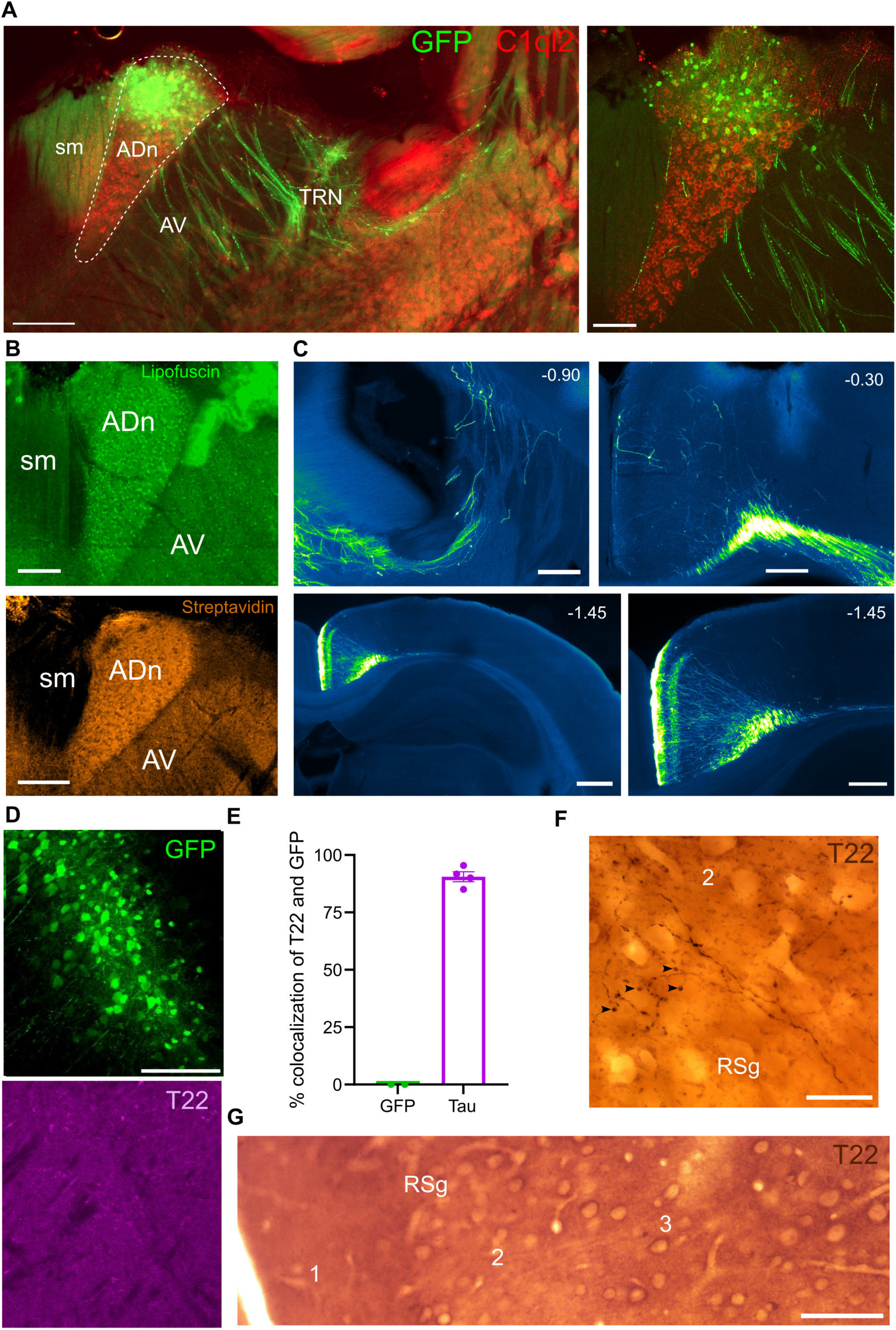
GFP expression and htau immunoreactivity in HD-tau and HD-gfp mice, related to. Figure 1. (**A**) Widefield epifluorescence showing localization of the ADn based on C1ql2 immunoreactivity (red, dashed region), case SJ14. Viral expression (GFP, green) was restricted to the dorsal part of the ADn. Axons of GFP-expressing ADn cells traverse the AV and branch in the TRN and course through the striatum. Right, enlarged view of ADn. (**B**) Top, the ADn has a high level of lipofuscin compared to surrounding brain regions (488 nm excitation widefield autofluorescence, case SJ14). Bottom, the ADn has a high density of biotin, visualized by streptavidin-cy5, compared to neighboring regions (widefield epifluorescence, case SJ25). (**C**) Trajectory of GFP-expressing ADn cell axons to postsynaptic target regions. Widefield epifluorescence micrographs, case SH30. Top left, axons leave the ADn and branch in the TRN at -0.90 mm from Bregma. Axons travel rostrally through the striatum and then turn caudally to enter the cingulum bundle (top right, -0.30 mm from Bregma). At the level of dorsal hippocampus (-1.45 mm from Bregma, bottom panels), ADn cell axons innervate the full extent of the RSg, then continue to other areas including PoS. (**D**) Lack of T22 immunoreactivity (magenta) in GFP-expressing cells (green) in the thalamus of an HD-gfp mouse (case SH30). Confocal maximum intensity z-projection (35 µm thick). (**E**) Quantification of GFP cells and T22-positive cells in HD-tau (case SH38, SH40, TV172, TV178) and HD-gfp mice (case SH49, SH56). (**F**) T22-immunoreactive axons and axon terminals (e.g. arrows) in RSg layer 2 (right) from an HD-tau mouse (case TV176). Light microscopic image (minimum intensity projection), DAB-based HRP reaction. (**G**) Lack of T22 immunoreactivity in the RSg from an HD-gfp mouse (case TV177). Light microscopic image, DAB-HRP reaction. Scale bars (µm): A left 200, A right 100; B 200; C top panels and bottom right 250, bottom left 500; D 100; E 20; F 50.

**Figure S2.**
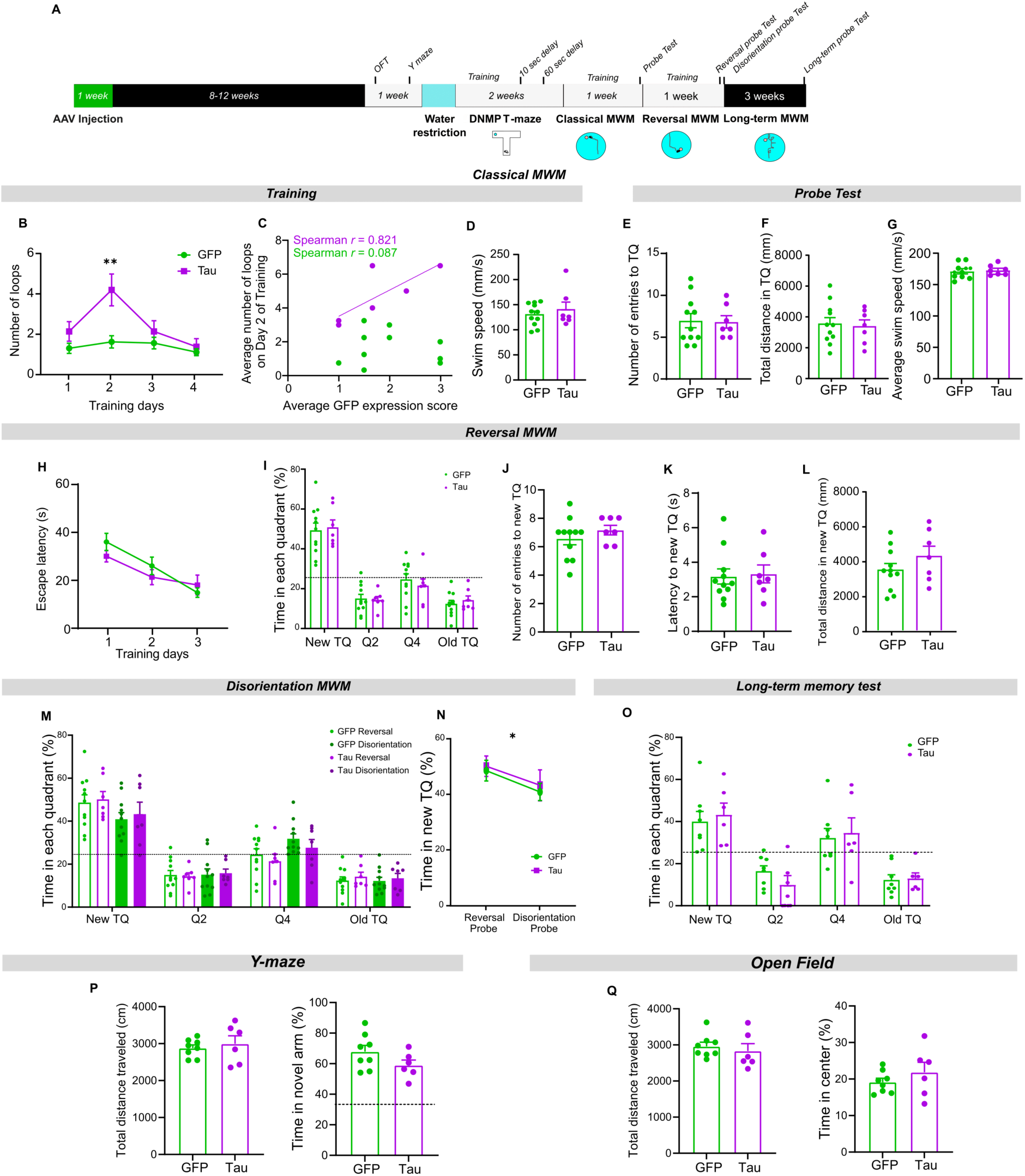
Behavioral tests in HD-tau and HD-gfp mice, related to Figures 2 and 3. (**A**) Experimental design; AAVs were injected into the ADn of mice. After 8-12 weeks of expression, mice were tested in a battery of behavioral tasks. (**B**) In the classical MWM, the average number of loops per training day was significantly different on day 2 (*group × training effect* two-way repeated measures ANOVA: *F*_3,48_ = 4.121, \**p* = 0.0111; *post-hoc* LSD test: \*\*\**p* <0.0001). (**C**) Correlation between the expression levels of GFP/htau-GFP and the average number of loops. (**D**) There was no difference in swim speed between groups on training day 2 (*t*_(16)_ = 0.7242, *p* = 0.4794). (**E-G**) During the MWM probe test, no differences were detected between groups for the number of entries to the TQ (E, (*t*_(16)_ = 0.1177, *p* = 0.9078), the total distance traveled in the TQ (F, *t*_(16)_ = 0.2664, *p* = 0.7934), or average swim speed (G, *t*_(16)_ = 0.291, *p* = 0.7748). (**H-L**) Both groups performed similarly during training in the reversal phase of the MWM (H, *training effect* two-way repeated measures ANOVA: *F*_2,32_ = 14.97, *\*\*\*p* < 0.0001), in the percentage of time spent in the TQ during the reversal probe test (I, HD-gfp vs chance*: U* = 6, \*\**p* = 0.0040; HD-tau vs chance: *U* = 6, \**p* = 0.0239, HD-gfp vs HD-tau: *t*_(16)_ = 0.4040, *p* = 0.6916), as well as in the number of entries to the TQ (J, *U* = 28.5, *p* = 0.3734), latency to new TQ (L, *t*_(16)_ = 0.2151, *p* = 0.8324), and the total distance traveled in the TQ (L, *t*_(16)_ = 1.260, *p* = 0.2258). (**M-N**) Proportion of time spent in each quadrant in the disorientation probe test (M). Note that both groups spent significantly less time in the new TQ after disorientation. Proportion of time spent in the new TQ in the reversal probe test versus in the disorientation probe test (N, *disorientation effect* two-way ANOVA: *F*_1,16_ = 5.240, *\*p* = 0.0360). (**O**) A long-term memory probe test was performed 3 weeks after the disorientation probe test. No differences were observed between the groups. Both groups showed a clear preference for the TQ (HD-gfp vs chance: \*\*\**p* = 0.0003; HD-tau vs chance: \*\*\**p* = 0.0006, HD-gfp vs HD-tau: *t*_(16)_ = 0.4381, *p* = 0.6691). (**P**) Spontaneous spatial novelty preference was tested using a Y-maze that consisted of three arms (start arm, S; other arm, O; novel arm, N). Bar charts for HD-gfp mice (n = 8) and HD-tau mice (n = 6) show no difference in the total distance traveled (left, *t*_(12)_ = 0.5559, *p* = 0.0626) or in the proportion of time spent in the novel arm (right, N/N+O, *t*_(12)_ = 1.592, *p* = 0.1374). (**Q**) No difference was observed between groups for the total distance traveled (*t*_(12)_ = 0.5889, *p* = 0.2913) or time spent in the center (*t*_(12)_ = 1.01, *p* = 0.0583) during the open field test. Dashed lines, chance.

**Figure S3.**
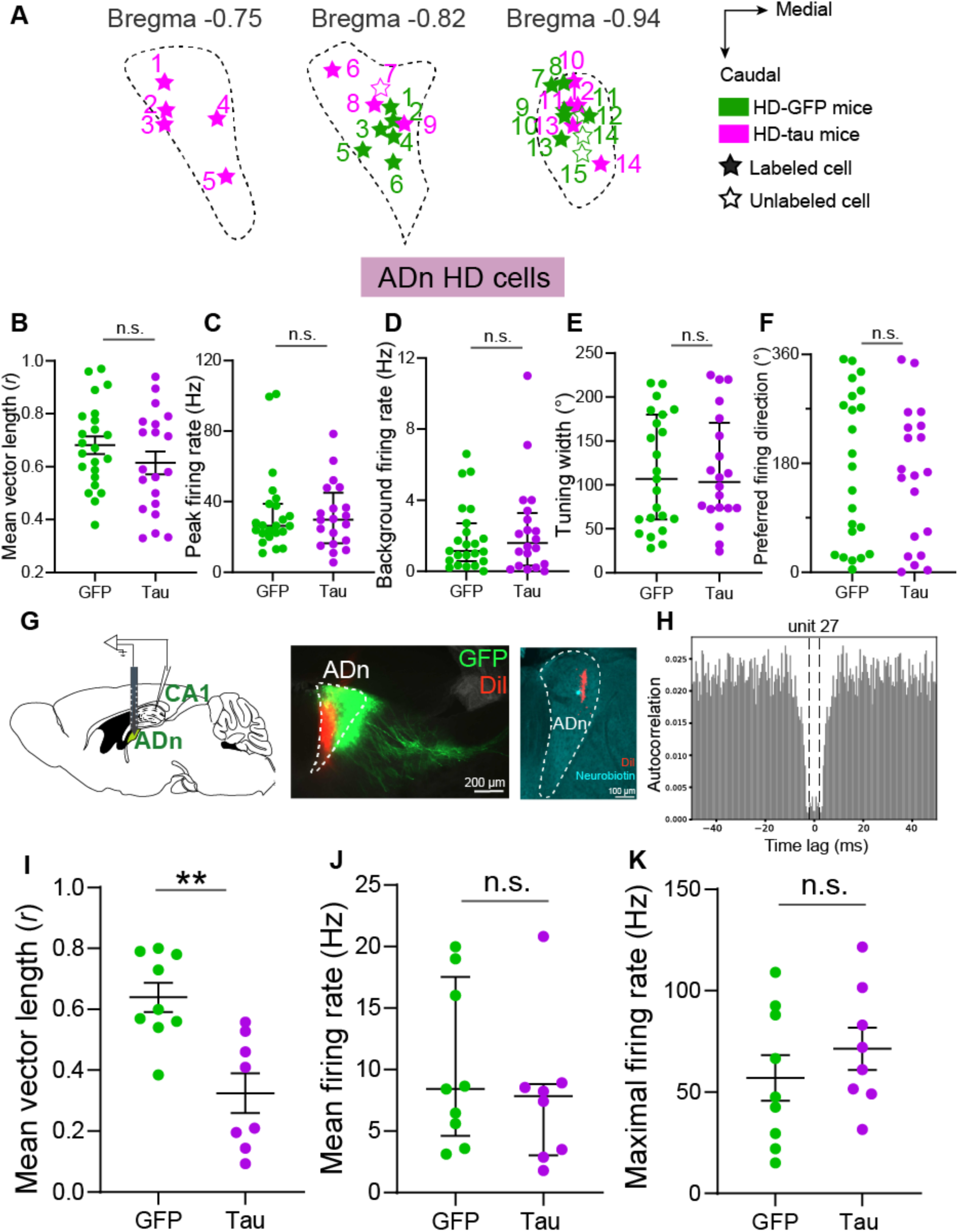
Recording locations and electrophysiological properties of ADn cells, related to Figure 4. (**A**) Schematic of the ADn at three representative anterior-to-posterior levels (coronal sections) indicated by the distance from Bregma in mm. Solid stars represent neurobiotin-labeled and identified ADn cells (n = 13 from 7 HD-tau mice, n = 12 from 5 HD-gfp mice). Unfilled stars represent unlabeled ADn cells recorded near to labeled cells (n = 1 from 1 HD-tau mouse, n = 3 from 1 HD-gfp mouse). Cell ID (from 1-14, HD-tau mice): TV189-17, TV189-16, TV189-15, SH37-03, SH36-2-h, SJ14-10, SH33-2-b, SH33-2-c, SJ16-10, SH40-061, SH40-08, SH40-071, SH40-062, SH40-05; cell ID (from 1-15, HD-gfp mice): TV190-2-h, SH46-2-d, SH30-10, TV190-2-i, SH56-2-a, TV191-3-f, TV191-3-e, SH49-021, SH49-022, SH49-023, SH46-2-e, SH56-2-c, SH49-031, SH46-2-f, SH46-2-g. (**B-F**) Directional properties of ADn HD cells recorded with glass electrodes (B, mean vector length; C, peak firing rate; D, background firing rate; E, directional tuning width; F, preferred firing direction). Peak firing rate: 2.99 [1.6-4.5] Hz from HD-tau mice versus 2.63 [2.2-3.8] Hz from HD-gfp mice, *U* = 223, *p* = 0.8708; background firing rate: 0.16 [0.03-0.32] Hz from HD-tau mice versus 0.12 [0.05-0.27] Hz from HD-gfp mice, *U* = 217, *p* = 0.7586; preferred firing direction: 411.5 ± 145.4° from HD-tau mice versus 363.9 ± 85.9° from HD-gfp mice, *U* = 200, *p* = 0.4765. (**G**) Left, schematic showing *in vivo* 16-channel probe recordings in the ADn. Middle, widefield fluorescence micrograph of a coronal brain section showing viral expression of GFP (green) and a probe tract (Dil, red) localized to the ADn, case SH46. Right, a coronal brain section showing a neurobiotin-labeled HD cell (SH33-2-c, cyan) and a probe tract (Dil, red) localized to the ADn. (**H**) The 100 ms autocorrelogram for all spikes from unit27 from mouse SH46. Bin size, 0.05 ms; window size, 50 ms; refractory period, 2 ms. (**I-K**) Mean vector length (I), mean firing rate (J, 7.8 [3.0-8.8] Hz for HD-tau mice versus 8.4 [4.6-17.5] Hz for HD-gfp mice, *U* = 29, *p* = 0.5414), and maximal firing rate (K, 71.4 ± 10.5 Hz for HD-tau mice versus 56.94 ± 11.22 Hz for HD-gfp mice, *t*_(15)_ = 0.9326, *p* = 0.3658) for sorted units from recordings with 16-channel probes in the ADn from HD-pau mice (Tau) and HD-gfp mice (GFP). \*\**p* < 0.01; not significant, n.s..

**Figure S4.**
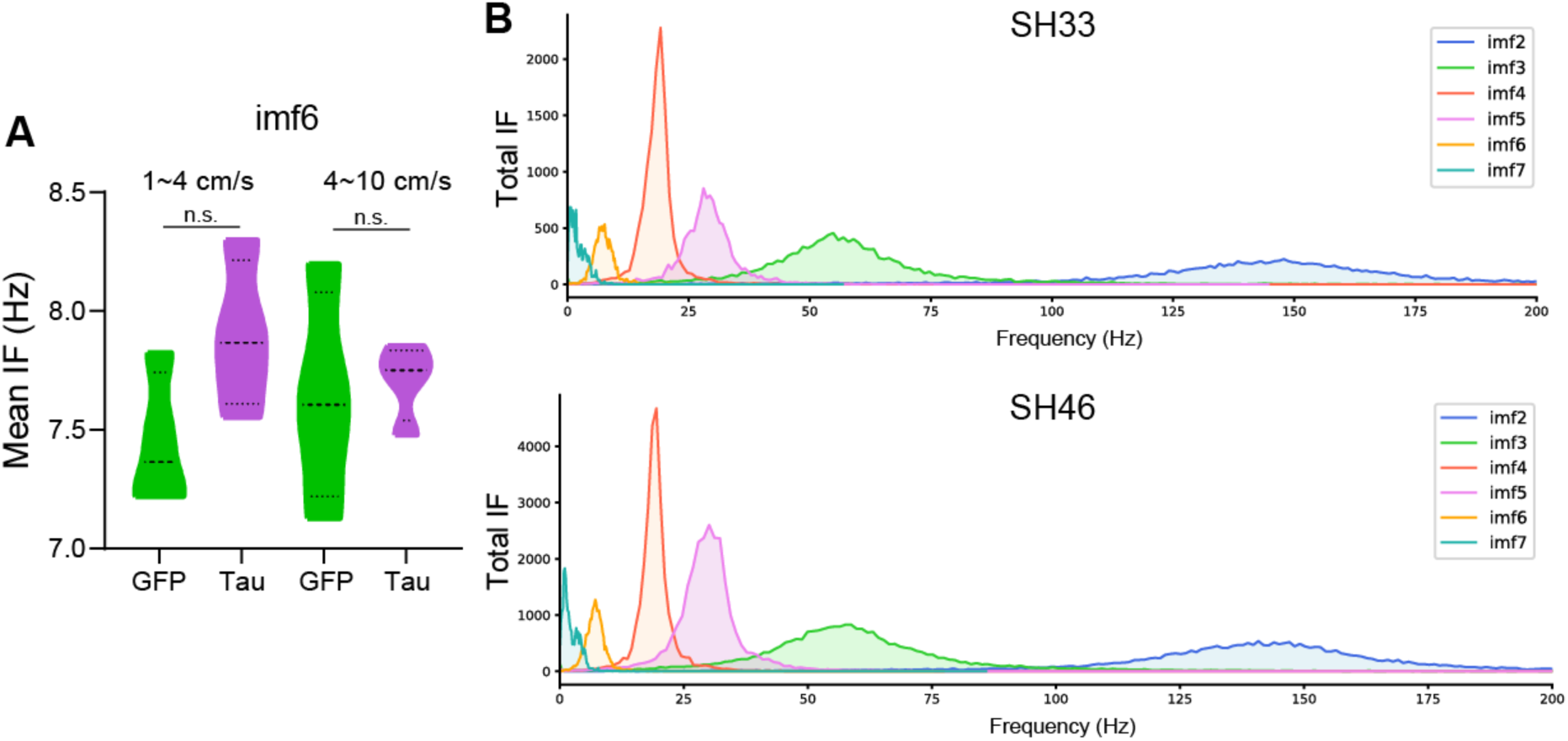
Unimpaired hippocampal theta oscillations in HD-tau mice, related to Figures 4 and 5. (**A**) Distribution chart of mean instantaneous frequency values in 4 HD-gfp mice (GFP, green) and 4 HD-tau mice (Tau, purple) for two speed bins (1–4 cm/s, *t*_(6)_ = 2.139, *p* = 0.0763; 4–10 cm/s, *t*_(6)_ = 0.3, *p* = 0.7688) during spontaneous movement on the running disc. (**B**) Instantaneous frequency for the intrinsic mode functions (IMFs, see Methods). Top, HD-tau mouse SH33. Bottom, HD-gfp mouse SH46. Abbreviations: IF, instantaneous frequency; n.s, not significant.

## Videos

Videos S1-5: The green circle represents the location of the hidden platform. Video speed: x2.

Videos S6-7: The green circle represents the previous location of the hidden platform during training. Video speed: x2.

**Video S1.** HD-gfp mouse SH31 during training on Day 2, Trial 2 of the MWM.

**Video S2.** HD-gfp mouse SJ26 during training on Day 2, Trial 2 of the MWM.

**Video S3.** HD-tau mouse SH38 during training on Day 2, Trial 3 of the MWM.

**Video S4.** HD-tau mouse SH33 during training on Day 2, Trial 2 of the MWM.

**Video S5.** HD-tau mouse SH40 during training on Day 2, Trial 2 of the MWM.

**Video S6.** HD-gfp mouse SH31 during probe of the MWM, following disorientation.

**Video S7.** HD-tau mouse SH40 during probe of the MWM, following disorientation.

## Resource Availability

### Lead contact

Further information and requests for resources and reagents should be directed to and will be fulfilled by the lead contact, Tim Viney.

### Materials availability

Apart from the viral vectors (see Key resources table), this study did not generate new unique reagents.

### Data and code availability

Code will be made available on GitHub.

## Acknowledgements

We thank Hugo Malagon-Vina for excellent advice on acute silicon probe data analysis and Violette Chiara for help with implementing the tracking software Animal TA. We also thank Mara Wülfing, Aditi Athreya, and Kathryn Holland for help with tissue processing. Funding: Alzheimer’s Society grant 522 AS-PhD-19a-010 (T.J.V., B.S.); Medical Research Council grants MR/R011567/1 and MR/Z504518/1 (T.J.V.); The John Fell Fund grant 0013781 (T.J.V.); UKRI grant EP/Z001358/1 (T.J.V., S.H.); NIH R01 MH120073 (M.E.H.); U.S. Office of Naval Research MURI N00014-1-19-2571 (M.E.H.); NIH F32MH139270 (P.A.L.). S.H. was supported by a Blaschko Fellowship from the Department of Pharmacology, Oxford. S.J. was supported by a Clarendon Scholarship.

## Author Contributions

Conceptualization: S.J., S.H., B.S., T.J.V. Methodology: S.J., S.H., B.S., V.G.G., D.B., T.J.V. Software: S.J., S.H., V.G.G., P.A.L. Validation: S.J., S.H., B.S., P.A.L., M.E.H., D.B., T.J.V. Formal analysis: S.J., S.H., B.S., V.G.G., T.J.V. Investigation: S.J., S.H., B.S., V.G.G., P.A.L., M.E.H., D.B., T.J.V. Resources: S.H., M.E.H., D.B., T.J.V. Data curation: S.J., S.H., T.J.V. Writing—original draft: S.J., S.H., T.J.V. Writing—review and editing: S.J., S.H., B.S., V.G.G., P.A.L., M.E.H., D.B., T.J.V. Visualization: S.J., S.H., B.S., V.G.G., T.J.V. Supervision: T.J.V. Project administration: T.J.V. Funding acquisition: S.H., T.J.V.

## Declaration of interests

The authors declare no competing interests.

## Methods

### Key resources table

| REAGENT or RESOURCE | SOURCE | IDENTIFIER |
| --- | --- | --- |
| **Antibodies** |  |  |
| T22 | Merck | Cat. No. ABN454 |
| HT7 | Pierce Thermo Scientific | Cat. No. MN-1000 |
| 4R Tau | Abcam | ab242333 |
| Tau (phospho T231) | Abcam | ab151559 |
| VGLUT2 | Synaptic Systems | Cat. No. 135 404, RRID:AB_887884 |
| Parvalbumin | Synaptic Systems | 195004/1-19, RRID: AB_2156476 |
| C1ql2 | Novus Biologicals | Cat. No.NBP2-34090 |
| **Chemicals, peptides, and recombinant proteins** |  |  |
| Ultra-purified recombinant AAV8 virus, pilot-scale packaging pAAV-CBh>EGFP:WPRE | Vector Builder | Catalog # AAV8SP(VB900088-2238xse)-C<br>Lot # 211102AAVJ09<br>https://en.vectorbuilder.com/vector/VB900088-2238xse.html |
| Ultra-purified recombinant AAV8 virus, pilot-scale packaging pAAV-CBh>EGFP:T2A:Tau(P301L):WPRE | Vector Builder | Catalog # AAV8SP(VB211015-1033pqq)-C<br>Lot # 211117AAVD11<br>https://en.vectorbuilder.com/vector/VB211015-1033pqq.html |
| Neurobiotin tracer | Vector Laboratories | #SP-1120; RRID: AB_2313575 |
| Isoflurane inhalation anesthesia | Teva Ltd. | https://products.tevauk.com/p/Category?id=185 |
| Vetergesic (0.3 mg/ml Buprenorphine) | Ceva<br>Marcain (Bupivacaine) | Aspen #PS09728 |
| Refobacin bone cement | Zimmer Biomet | #3003940002-3 |
| Tetric EvoFlow dental cement | Ivoclar Vivadent | A1 |
| Dil tracer | Life Technologies | #V22889 |
| Normal horse serum | Vector Laboratories | S-2000 |
| Normal goat serum | Vector Laboratories | S-1000 |
| **Critical commercial assays** |  |  |
| Vectastain ABC Elite kit | Vector Laboratories | Cat# PK6100; RRID: AB_2336819 |
| Vectashield Antifade Mounting Medium | Vector Laboratories | Cat# H-1000; RRID: AB_2336789 |
| **Experimental models: Organisms/strains** |  |  |
| C57BL6j mice | Charles River Laboratories | https://www.criver.com/ |
| **Software and algorithms** |  |  |
| FIJI (ImageJ) | NIH | https://imagej.net/software/fiji/ |
| MATLAB | Mathworks | https://www.mathworks.com/products/matlab.html |
| Python |  | https://www.python.org/ |
| Zen Black, Zen Blue, Axiovision | Zeiss | www.zeiss.co.uk |
| Spike2 | Cambridge Electronic Design | http://ced.co.uk/ |
| Kilosort2 | (Pachitariu et al., 2024) | https://github.com/MouseLand/Kilosort |
| Phy |  | https://github.com/cortex-lab/phy |
| GraphPad Prism 7 | GraphPad Software | https://www.graphpad.com/ |
| AnimalTA | (Chiara and Kim, 2023) | http://vchiara.eu/index.php/animalTA |
| **Other** |  |  |
| A1x16-poly 2s silicon probe | NeuroNexus | https://neuronexus.com/ |

### Experimental model and subject details

A total of 53 adult (29 female) C57Bl6j mice (obtained from Charles River Ltd, UK) were injected with adeno-associated viruses (AAVs) targeted to the anterior thalamus. Inclusion/exclusion criteria were as follows: n = 41/53 mice had viral expression in the ADn and were used for histology; n = 12/53 mice were included in pilot behavioral tests; n = 36/53 mice were included in the main behavioral tests (Fig. 2, 3, S2), n = 11/36 were excluded due to a lack of sufficient viral expression in the ADn; n = 32/53 mice were included for *in vivo* recordings to screen for HD cells, n = 2/53 were used for *in vitro* recordings (not reported), n = 11/32 mice were excluded due to a lack of viral expression in the recording locations; 26 mice that were included in behavioral tests and/or *in vivo* recordings are reported in Table 1. Recordings from excluded mice will be reported elsewhere.

All animal procedures were approved by the local Animal Welfare and Ethical Review Body under approved personal and project licenses in accordance with the Animals (Scientific Procedures) Act, 1986 (UK) and associated regulations. Animals were maintained in individually ventilated cages on a 12/12 h light-dark cycle (lights on at 07:00). They were typically group-housed (4–5 per cage) to prevent detrimental effects of social isolation. Individual cages contained mice from both groups with same sex. Room temperature was maintained at 21°C and humidity was between 50% and 60%. Mice were fed on a standard RM3 diet (pellets, product 801700, Special Diets Services).

### Surgical procedures

#### Viral tracing

Mice (5-9 weeks old) were anesthetized with isoflurane, the scalp was clipped, and a subcutaneous injection of Buprenorphine (0.1 mg/kg dose, Vetergesic, Ceva) was administered. Mice were fixed to a stereotaxic frame via ear bars and a jaw bar. Anesthesia was maintained with 1–3% (v/v) isoflurane and body temperature was regulated with a homeothermic blanket (Harvard Apparatus). Ocular lubricant was applied to the eyes, and Bupivacaine (Marcaine, Aspen) was injected into the scalp. An incision was made along the scalp to expose the skull. Craniotomies were made using a surgical microdrill at the following coordinates to target the ADn bilaterally (mm from bregma): -0.85 antero-posterior (AP), ± 0.75 medio-lateral (ML). A glass pipette was gradually lowered to 2.7 mm below the brain surface and 100-200 nl of adeno-associated virus (AAV) was pressure injected, either Tau AAV (pAAV-CBh>EGFP:T2A:Tau(P301L):WPRE, 1.335 x 10^13^ GC/ml) or control GFP AAV (pAAV-CBh>EGFP:WPRE, 5.9 x 10^12^ GC/ml), and left in place for 5 minutes before withdrawal. After the injections, the scalp was sutured and animals recovered in a cage over a heated blanket.

### Behavioral tests

All behavioral tests were carried out during the daytime. Mice were first handled for 3 days before initiating behavioral testing and were subjected to a battery of behavioral tests following the order in Fig. S2A: open field test (Day 1), Y maze (Day 2), T-maze (Days 8-23, following 3 days of water restriction and 2 days of habituation) and MWM (Days 24-34, 48). All tests except for the T-maze and MWM were performed as previously described (Viney et al., 2022). Behavioral tests were recorded using a camera (Microsoft LifeCam, 30 frames/s), and parameters were extracted and analyzed using AnimalTA software (Chiara and Kim, 2023). Some animals were excluded from the analysis due to technical issues with the video recordings (Fig. 3E,F, S2P,Q).

#### Open field test

Mice were placed individually in the center of an open field chamber (consisting of an open 50 x 50 cm white square arena and 30 cm high black plastic walls) for 5 minutes. The room was illuminated using lamps in the ceiling above the arena (340-460 lux light intensity). The arena was thoroughly cleaned with scentless Anistel between subjects. The center area was defined as the 25 × 25 cm interior portion of the square arena. Each mouse was placed gently in one corner of the arena. The total distance traveled (cm) and time spent in the center area were quantified.

#### Spatial novelty preference Y-maze

The task was designed to test spontaneous exploration and the innate preference for exploring a novel environment, as previously reported (Sanderson et al., 2007; Viney et al., 2022). A solid black plastic Y-maze (consisting of three 30 cm x 7 cm arms with 20 cm high walls, 160-320 lux light intensity) was positioned 17 cm above the floor. The maze had no intra-maze cues, except for a 20 cm high magenta block that obstructed one of the arms during the sample phase. Prominent extra-maze cues were present.

The test was divided into an *exposure phase* and a *test phase*. During the exposure phase, the novel arm was blocked for 5 minutes, and the mice were placed at the end of the start arm, allowing them to explore the start and other arms. In the test phase, the block was removed when the mice were facing away from it, and they were then free to explore all three arms for an additional 5 minutes. To prevent the influence of odor cues, the maze was cleaned with Anistel after each mouse’s visit. Time spent and distance traveled in each arm were analyzed. An entry into an arm was registered when the center of the mouse’s body entered the defined arm area.

#### Non-matching-to-place T-maze

Spatial working memory was assessed as previously reported (Roy et al., 2021; Viney et al., 2022). Mice were water-restricted until they reached 85-90% of their initial body weight (around 3 days). During this time, mice were habituated to the water reward, which would subsequently be used as a reward in the non-matching-to-place T-maze task. Before the training phase, mice were habituated for 10 min to the T-maze apparatus (50 x 10 cm start arm and two identical 30 x 10 cm goal arms; walls were 10 cm high, and the maze was raised 44 cm off the floor, with a light intensity of 330-520 lux). The rewards consisted of 0.5 ml of water in small plastic dishes, placed at the ends of both goal arms. The habituation was done first in groups (with cagemates) and then individually.

The habituation stage was followed the next day by the *training stage,* which consisted of 4 trials per day (defined as a *session*) with each trial having two separate phases (sample and choice phases). In the sample phase, the presence of a block forced the mouse to visit one arm where it was free to retrieve its reward (0.1 ml of water). After reward consumption, mice were returned to their home cage for around 15 s and the T-maze was quickly cleaned. This was followed by the choice phase in which the block was removed and the mouse was free to visit both arms: either visiting the previously unvisited arm to obtain a second 0.1 ml reward or revisiting the previous arm, which was now unrewarded. A pseudorandom sequence was used to assign the rewarded arm, with the same numbers of left or right turns per session.

Following 10 days of training, a *delay stage* was introduced. Mice were blocked at the starting point of the maze between the sample and choice phases for either 10 s (8 trials over 2 days) or 60 s (16 trials over 4 days) with inter-trial intervals of ∼4–8 min, depending on cage size. The *alternation rate* was calculated as the average score across trials: a score of 1 was given if the mouse chose the rewarded arm, and a score of 0 if it chose the unrewarded arm. The *decision latency* (the time duration from when the mouse starts to move to when it makes a decision, i.e. enters one of the goal arms) and the *average number of head turns* per session (the number of head turns during the time from when the mouse starts to move to when it makes a decision) were quantified for choice runs from the 60 s delay sessions (4 trials). Head turns, defined as rotations of the head away from or back to the midline position of the body along the azimuthal plane, were manually counted by an observer blinded to the group assignments.

#### Morris water maze (MWM)

Following two days of recovery from the water restriction protocol, mice were trained on the *classical MWM protocol*. Mice were placed facing the sidewall of a plastic pool (120 cm diameter, 30 cm high walls, 30-100 lux light intensity) filled with water (15 cm depth, ∼22℃) and containing a hidden platform (10 cm diameter, 14 cm high) submerged in the water. Each training day consisted of four trials during which mice were placed at four different locations of the pool (N, W, E, or S) as the start points. The order of the start points was randomized each day for both groups. The hidden platform was always located in the quadrant between N and W (defined as the target quadrant). Mice were given 60 s to find the hidden platform. In the event of an unsuccessful search, mice were gently guided to the hidden platform where they remained for around 10 s. The next trial was then initiated. Mice were placed back in their home cage after 2 trials for an inter-trial period of ∼2 min before completing trials 3 and 4. Mice were thoroughly dried, and cages were continuously placed on a heat-pad to keep the mice warm between trials. The escape latency during training was assessed as the average time spent to find the hidden platform across the four trials. All trials were recorded via a video camera, and the swim path was quantified using AnimalTA. All experimenters were blind to the experimental group.

The strategy score was quantified manually based on a previously described method (Berkowitz et al., 2018). Using the tracking data obtained from AnimalTA software, we conducted a detailed qualitative analysis of swim trajectories to assess the number of directed swim movements toward the hidden platform during training. One experimenter manually scored these movements based on definitions established in prior research (Garthe et al., 2009; Gehring et al., 2015-10-01; Berkowitz et al., 2018), blind to the group. Mice were scored from 1 to 4 depending on their favored search strategy before finding the platform on each trial: 1 = target-direct strategy (direct path, circuitous direct path with no loops), 2 = target-indirect strategy (circuitous indirect path), 3 = spatial-indirect strategy (chaining, scanning, focused search), 4 = non-spatial strategy (incursion, looping, thigmotaxis). Loops, defined as a mouse swimming in a small circle along a swim path, was manually counted for every trial.

On the 5^th^ day, the hidden platform was removed and mice were released at the opposite quadrant to the normal platform location, facing the wall for the probe test (P1). The trial lasted for 60 s and the time spent and distance swam in each quadrant of the pool were quantified. Other parameters that were extracted were as follows: latency to reach the TQ, the number of entries to the TQ. The swim speed and total distance traveled were also quantified.

Next, *reversal training* was carried out, which consisted of 3 days of training with 4 trials per day. The hidden platform was placed in the opposite quadrant (new TQ) from the original location. After 3 days of reversal training, the hidden platform was removed and another probe test was performed (P2). The same parameters obtained for the classical MWM were extracted for the reversal MWM. Following the reversal probe test, mice from both groups were disoriented by being placed in a small dark box and rotated 16 times in alternating back-and-forth directions at a speed of 180° to 270°/s continuously for 60 s in the same room that contained the MWM(Grieves et al., 2022-11-01). Mice were then immediately placed in the pool for another probe test (‘disorientation probe test’, P3). Finally, three weeks after P3, mice underwent a final probe test (P4) to examine long-term memory.

### *In vivo* electrophysiology

#### Habituation

After completing all behavioral tests, mice were handled for 4 days prior to headplate implant surgeries and were habituated to the recording room by being placed on the running disc setup and being free to explore the apparatus at least 5 min per day (Fig. 4A).

#### Headplate implant and craniotomies

Mice were prepared as above for the *viral tracing* surgery. After exposing the skull, two M1 screws (Precision Technology Supplies) were fixed into the skull above the cerebellum as anchor points, with one or both used as electrical reference. Another screw was fixed ∼1.50 mm anterior of bregma. Screws were sealed with Refobacin bone cement (Zimmer Biomet), and a machined glass-reinforced plastic D-shaped headplate (custom made at the Department of Physics, Oxford University) was positioned over the screws and secured with bone cement. Mice were administered a subcutaneous injection of 0.5 ml 5% (w/v) glucose in saline (Aqupharm 3, Animalcare) peri-operatively. Craniotomies were made using a surgical microdrill at the following coordinates (mm from bregma): Hippocampal CA1, -2.5 AP, ± 1.5 ML; ADn -0.85 AP, ± 0.75 ML. The dura was removed with a bent 27 gauge needle. Silicon (Smooth-On) was applied to protect the craniotomy sites by covering the skull inside the headplate, and mice recovered in a cage over a heated blanket. Electrophysiology experiments were initiated the following day.

#### Extracellular recordings with glass electrodes

Mice were head-restrained via a custom-made stainless-steel block (Department of Physics, Oxford University) secured to a heavy-duty frame (model 1430, Kopf Instruments). Head-restrained mice could spontaneously run and rest on a 30 cm diameter plastic running disc covered with paper roll (Fig. 4A). The apparatus was mounted to a turntable consisting of a circular aluminum breadboard (Thorlabs Inc) enabling 360° passive rotation. Recordings were performed under photopic conditions (600-700 Lux). Two glass electrodes filled with 3 % neurobiotin (w/v) in 0.5 M NaCl (8–18 MΩ) were lowered via the craniotomy sites using IVM-1000 micromanipulators (Scientifica Ltd) to reach the hippocampus and ADn. Signals were amplified x1000 (ELC-01MX and DPA-2FS modules; npi Electronic GmbH, Tamm, Germany). Wide-band (0.3 Hz to 8 kHz) and band-pass-filtered (action potentials, 0.8–8 kHz) signals were acquired and digitized at 20 kHz (Power1401; Cambridge Electronic Design Ltd, Cambridge, UK). HumBugs (Digitimer Ltd) were used to remove 50 Hz noise. Speed was measured using a rotary encoder (HEDM-5500#B13, Avago Technologies) underneath the running disc. An inertial measurement unit (SparkFun OpenLog Artemis, which is preprogrammed to automatically log data from Global Navigation Satellite System navigation data) attached to the turntable enabled monitoring HD data at 10 Hz. Electrophysiology data were recorded using Spike2 software (Cambridge Electronic Design). Local field potentials (LFPs) were recorded from the stratum pyramidale of the CA1 region of the hippocampus, identified by the combination of positive sharp-waves and 130–230 Hz ripples during rest periods and 5–12 Hz theta oscillations during movement periods (Viney et al., 2022). The ADn was targeted bilaterally in each mouse. To localize the ADn, the glass electrode was lowered through the craniotomy sites to the target depth (∼ 2.5 mm). HD cells were detected by slowly rotating the apparatus until an abrupt increase in firing was observed when the mouse faced a specific direction. The firing activity was then recorded while the mouse was manually rotated both clockwise and counterclockwise to acquire firing data across all angles. Cells of interest were then juxtacellularly labeled with 200 ms positive current pulses, followed by a recovery period of 2-6 h.

#### Acute silicon probe recordings

Following glass electrode recordings and localization of the ADn, methylene blue was applied to the tip of the glass electrode and lowered to the same AP-ML position in the craniotomy site to ensure proper localization of the probe. Then an acute 16-channel silicon probe (A1x16-poly 2s probe, NeuroNexus) connected to an RA16-AC preamplifier (Tucker-Davis Technologies) was carefully painted with Dil and positioned at the exact location marked by the methylene blue tracer then lowered to the same depth as the previous glass electrode. Signals were amplified x1000 (Lynx-8 amplifiers, Neuralynx), band-pass filtered (0.5 Hz to 8 kHz), digitized at 20 kHz with a Power1401 and recorded with Spike2. Silicon probe recording sites were confirmed by examining Dil fluorescence in serially processed brain sections.

### Histology

#### Transcardial perfusion and sectioning

Mice were deeply anesthetized with sodium pentobarbital (50 mg/kg, i.p.) and transcardially perfused first with saline followed then with 4% paraformaldehyde, 15% v/v saturated picric acid, and 0.05% glutaraldehyde in 0.1 M phosphate buffer (PB), pH 7.4. Brains were removed from the skull. Some brains were post-fixed overnight in fixative lacking the glutaraldehyde. After washing in 0.1 M PB, 70 µm coronal sections were cut using a Leica Microsystems VT 1000S vibratome and stored in 0.1 M PB with 0.05% sodium azide at 4°C.

#### Streptavidin visualization and immunohistochemistry

Using the indirect primary antibody detection method in combination with fluorochrome-conjugated secondary antibodies, the expression of cell-type specific molecules was tested on individually labeled neurons and on control tissue.

For the visualization of neurobiotin-labeled cell processes, brain sections were permeabilized in Tris-buffered saline (0.9% NaCl buffered with 50 mM Tris, pH 7.4; TBS) with 0.3% Triton X-100 (TBS-Tx) or via rapid 2x freeze–thaw (FT) over liquid nitrogen (cryoprotected in 20% sucrose in 0.1 M PB) then streptavidin-conjugated Cy3 or Cy5 was applied at 1:500 dilution in TBS-Tx (or TBS if permeabilized with FT) for 4 h at room temperature (RT) or overnight at 4°C. Sections were washed in TBS/TBS-Tx and mounted to glass slides in Vectashield (Vector Laboratories).

First, sections were blocked for 1 h in 20 % normal horse serum (NHS) in TBS/TBS-Tx followed by 2-6 days incubation at 4°C in primary antibody solution containing 1 % NHS in TBS/TBS-Tx. The following primary antibodies (and dilutions) were used: rabbit anti-T22 1:1000 (ABN454, Merck), mouse anti-HT7 1:1000 (MN-1000, Pierce Thermo Scientific), rabbit anti-4R tau 1:1000 (ab242333, abcam), rabbit anti-Tau (phospho T231) 1:1000 (ab151559, Abcam), rabbit anti-C1ql2 1:1000 (NBP2-34090, Novus Biologicals), guinea pig anti-VGLUT2 1:1000 (135 404, Synaptic Systems), guinea pig anti-Parvalbumin 1:1000 (195004/1-19, Synaptic Systems). Subsequently, sections were washed 3 times in TBS/TBS-Tx, then incubated in secondary antibody solution containing 1 % NHS in TBS/TBS-Tx for 4 h RT or overnight at 4°C. The following secondary antibodies (and dilutions) were used in various combinations (all raised in donkey): anti-mouse and goat anti-rabbit Alexa Fluor 405 1:250 from Invitrogen, anti-guinea pig DyLight 405 (706-475-148), anti-guinea pig, anti-rabbit, and anti-mouse Alexa Fluor 647 1:500 (706-475-148, 711-605-152, 705-605-151) from Jackson ImmunoResearch. After 3 washes, sections were mounted in Vectashield.

For diaminobenzidine (DAB)-based horseradish peroxidase (HRP) reactions, sections were blocked for 1 h in 20 % normal goat serum (NGS) in TBS followed by incubation with primary antibodies in TBS containing 1 % NGS for 2 d at 4°C. After incubation with primary antibodies, sections were rinsed three times in TBS and blocked for 10 min at RT in 1% H_2_O_2_ solution in 0.1 M PB to reduce non-specific background reactions. Next, sections were incubated overnight at 4°C in 1:100 biotinylated goat anti-rabbit IgG (Vector Labs, BA-1000) in TBS containing 1% NGS. After repeated washes in TBS, sections were incubated for 3 days at 4°C in avidin-biotinylated horseradish peroxidase complex (Vectastain ABC Elite kit, Vector Laboratories) in TBS. Subsequently, peroxidase was visualized using a mix of 1% nickel ammonium sulphate, 0.4% ammonium-chloride, and 3,3-DAB (0.5 mg/ml, Sigma-Aldrich) developed with 0.01% H_2_O_2_. After washing sections in PB, sections were treated with 0.25% OsO_4_ in 0.1 M PB for 5 min. Next, after washing in 0.1 M PB at least four times, some sections were transferred onto slides in chrome alum gelatin and dried in air. To avoid air bubbles, sections were then incubated in fresh xylene for 10 min before being quickly mounted in DePeX mounting medium. Other sections were dehydrated using an ascending ethanol series followed by acetonitrile. This was followed by embedding in epoxy resin (Durpucan AMC, Fluka, Sigma-Aldrich), incubating overnight at RT then transferring onto slides. For polymerization, sections were incubated at 60 ℃ for 2 d.

### Microscopy

#### Wide-field epifluorescence- and confocal laser scanning microscopy

The immunohistochemical reactions were first evaluated using widefield epifluorescence on a Leitz DMRB microscope (Leica Microsystems) equipped with PL Fluotar objectives (magnification/numerical aperture: 5x/0.15, 10x/0.3, 20x/0.5, 40x/0.7; OpenLab software) or an AXIO Observer Z1 microscope (LSM 710; Zeiss) equipped with Plan-Apochromat 10x/0.3, 20x/0.8 and 40x/1.4 objectives (Axiovision or ZEN Blue 2.6 software). To improve image resolution and increase contrast, when needed, reactions were further assessed using confocal laser scanning microscopy; the LSM 710 was used with Plan-Apochromat 40x/1.4, 63x/1.4, and 100x/1.46 objectives (ZEN Black 14.0 software). Laser lines (solid-state 405 nm, argon 488 nm, HeNe 543 nm and HeNe 633nm) were configured with the appropriate beamsplitters. The pinhole was set to ∼1 Airy Unit for each channel.

### Code and Analysis

#### Classification of ADn and HD cells

Neurons that exhibited bursting non-rhythmic firing that was clearly related to the animal’s head direction were initially selected for recording, which biased the initial search strategy to HD cells of the ADn. By visualizing labeled cells, we confirmed that recorded cells from either the same penetration site or from closely aligned coordinates were located in the ADn. For glass electrode single-cell data, spikes were isolated by thresholding high-pass filtered voltage traces of their peaks and validated by using principal component analysis and/or visual inspection in Spike2 software (Cambridge Electronic Design, Cambridge, UK).

Analysis procedures were applied to both glass electrode single-cell data and silicon-probe data. Inclusions criteria: we analyzed recordings that were longer than 100 s, were from recording sites in the ADn that had GFP expression (Table 1), and had head direction coverage of at least one full turn with all angles represented. The directional tuning curve for each cell was obtained by plotting the firing rate as a function of the mouse’s directional heading, divided into bins of 6°. The firing rate was computed based on the total number of spikes divided by the total time in that bin. All HD properties-related parameters were analyzed based on published parameters (Taube, 1995) using customized Python scripts. The mean vector length (*r*) is a measure of the non-uniformity (or directionality) of the directional tuning curve and can vary between 0 (a uniform distribution) and 1 (a non-uniform distribution). Mean vector lengths were used with Rayleigh’s test to determine the extent of uniformity in the distribution of firing rate bins. Based on the published criteria (Peyrache et al., 2015; Blanco-Hernández et al., 2024), subjective assessment of the distribution of *r*, and each cell’s directional tuning curves, a criterion of *r* > 0.3, probability of non-uniform distribution < 0.001, and peak firing rate > 1 Hz was used to classify HD cells. For some cells and units that only showed directionality within a specific rotation direction (either *r*_CW_ (clockwise) or *r*_CCW_ (counterclockwise) > 0.3 and |*r*_CW_-*r*_CCW_| > 0.2), only spikes or units recorded during the most directionally responsive rotation direction were analyzed. Further details of HD cells exhibiting direction selectivity will be reported elsewhere.

The directional tuning curve was computed and fitted with von Mises distributions (Blair et al., 1997; Peyrache et al., 2015) or the best-fit triangular function (Taube et al., 1990) to avoid deviations caused by small fluctuations in the raw directional tuning curve. Four parameters were analyzed: (1) the preferred firing direction – the head direction angle with the highest firing rate; (2) peak firing rate – the highest firing rate from the directional tuning curve, which indicates the firing rate when the mouse is facing in the cell’s preferred direction; (3) directional tuning width – the range of head directions over which the cell fires, defined as 6x the s.d. to match the best-fit triangular function (Taube et al., 1990); and (4) background firing rate – the average firing rate when the mouse is facing outside of the directional firing range of the cell.

Another plot showing the number of spikes versus time was constructed to compute two parameters: (1) the maximum firing rate, which is defined as the highest firing rate within a 200 ms time bin from the entire recording session; (2) the mean firing rate, which is the average firing rate of the cell over the entire recording session.

#### Burst analysis

Burst analysis was performed using a burst analysis script in Spike2 (https://ced.co.uk/downloads/scriptspkanal). For each HD cell, spikes were extracted from time periods where the mouse was resting and facing the preferred firing direction. The minimum duration of the recording was 20 s, and the minimum detected bursts were set to 4. Recordings that did not meet these thresholds were excluded. The maximum initial interval signifying burst onset was set to 0.05 s and the longest inter-spike interval allowed within a burst was set to 0.03 s. The following parameters were extracted: percentage of spikes per burst (number of spikes within bursts/number of all spikes); the mean number of spikes per burst (average number of spikes within bursts); the mean burst length (average duration of bursts); the inter-spike interval during burst (duration between spikes during a burst); the inter-burst duration (duration between two consecutive bursts); and the mean cycle period (average period between bursts, cycle period = inter-burst duration + burst length).

#### Silicon probe data

Similar to the analysis of glass electrode recorded cells, only recordings from a Dil track confirmed to be within the ADn along with GFP expression and whose head direction coverage included at least one full turn were included in the analysis (Table 1). Spikes were detected, sorted, and clustered offline using Kilosort2 (Pachitariu et al., 2024) in MATLAB. Subsequently, interactive visualization and manual curation of the data were carried out using Phy2 based on refractory periods, spike waveform and cross-correlations. In order to obtain well-isolated and stable units, only units with mean firing rates > 1 Hz, refractory-period contamination < 2 ms, and consistent spike waveforms but dissimilar to nearby clusters were included in the analysis.

#### LFP analysis

LFP recordings with a regular synchronized theta rhythm and non-theta epochs containing sharp wave ripples were selected for LFP analysis. For LFP channels, direct current shifts were removed with 0.3 s sliding windows then channels were downsampled to 1.25 kHz in

Spike2 and were exported for analysis using customized Python scripts. 1 to 5 theta epochs were sampled per recording for each animal, ranging from 8 to 52 s to define each epoch. Values obtained per mouse were binned and averaged for low running speeds (1–4 cm/s) and ‘high’ speeds (4–10 cm/s). Empirical mode decomposition (EMD) was carried out in Python using the emd package (https://gitlab.com/emd-dev/emd) (Quinn et al., 2021). To obtain intrinsic mode functions (IMFs), the following masks were used for masked EMD (emd.sift.mask_sift): 350, 200, 70, 40, 30, 7, 1, divided by the sampling rate. Instantaneous amplitude, frequency and phase were obtained for each IMF using emd.spectra.frequency_transform with the Normalized-Hilbert Transform (nht). We excluded IMF1 as it contained mostly noise and excluded any mid-gamma range IMF that had 50 Hz electrical noise or poor movement periods (low signal to noise ratio).

### Quantification and statistical analysis

All data are represented as mean ± SEM or median [IQR]. Experimental units (e.g. mice, cells) are specified in the text after the n values. Statistical analysis was carried out in GraphPad Prism and Python. The alpha was set to 0.05. For data that approximated a normal distribution (tested by the Shapiro-Wilk test), unpaired Student’s t tests were used to compare two groups with equal variances and unpaired t tests with Welch’s correction were used in two groups having different variances, otherwise Mann-Whitney tests were used. For comparisons of more than two groups we used Analysis of Variance (ANOVA) for parametric data followed by Fisher’s LSD *post-hoc* test. For analysis of circular data, we used the Rayleigh test. Spearman’s correlation was used to test the correlation between two ranked variables.

## Notes

### Competing Interest Statement

The authors have declared no competing interest.

### Summary of Updates

Made some minor revisions to Figures 1, 4, S1, S3. Updated the Introduction, included some minor updates to the reporting of Results, and also updated parts of the Discussion.

